# A reference single-cell regulomic and transcriptomic map of cynomolgus monkeys

**DOI:** 10.1101/2022.01.22.477221

**Authors:** Jiao Qu, Fa Yang, Tao Zhu, Yingshuo Wang, Wen Fang, Yan Ding, Xue Zhao, Xianjia Qi, Qiangmin Xie, Qiang Xu, Yicheng Xie, Yang Sun, Dijun Chen

## Abstract

Non-human primates (NHP) are attractive laboratory animal models that accurately reflect both developmental and pathological features of humans. Here we present a compendium of cell types from the cynomolgus monkey *Macaca fascicularis* (denoted as ‘Monkey Atlas’) using both single-cell chromatin accessibility (scATAC-seq) and RNA sequencing (scRNA-seq) data at the organism-wide level. The integrated cell map enables in-depth dissection and comparison of molecular dynamics, cell-type composition and cellular heterogeneity across multiple tissues and organs. Using single-cell transcriptomic data, we inferred pseudotime cell trajectories and cell-cell communications to uncover key molecular signatures underlying their cellular processes. Furthermore, we identified various cell-specific *cis*-regulatory elements and constructed organ-specific gene regulatory networks at the single-cell level. Finally, we performed a comparative analysis of single-cell landscapes among mouse, cynomolgus monkey and human, and we showed that cynomolgus monkey has significantly higher degree of cell-type similarity to human than mouse. Taken together, our study provides a valuable resource for NHP cell biology.

## Introduction

Non-human primates (NHP) are phylogenetically close to humans and show various human-like characteristics, including genetics, organ development, physiological function, pathological response and biochemical metabolism. Hence NHP are extremely valuable as experimental animal models in medical research and drug development^1^. Since cells are the fundamental unit of all life, direct comparison of cell identities and cell-type compositions between organisms across organs would help to transfer knowledge in primates to medical research. In this regard, it is of vital importance to understand the cellular composition and heterogeneity of primate organs.

Rapid advances in single-cell multi-omics technologies have enabled molecular quantification of thousands of cells at once, leading to meticulous insight into organ compositions and mechanisms driving cellular heterogeneity^2^. Previous studies^3–7^ have mapped the single-cell landscapes across multiple organs in humans and mice, expanding our knowledge about the cellular heterogeneity underlying normal development and aging. Three-dimensional multicellular culture systems combined with single-cell transcriptome sequencing technology enables to chart the cellular and molecular dynamic changes during organ growth and development^8–10^. In addition, extensive efforts^9–12^ have been achieved to investigate how cells are perturbed in various disease conditions, including cancer and neurological disorders.

Mice have long been used as a representative model organism for mammalian development and physiology. Recently, extensive comparative analyses based on single-cell transcriptomics data have shown that both cell types and associated gene regulatory networks are conserved between human and mouse^13–20^, which provides a new perspective for explaining disease mechanisms and finding targets for disease intervention. However, it has been widely recognized that there are significant differences between mice and humans in terms of development and physiology^21^. From genetic perspective, primate experiments are more useful as they can better simulate human diseases and promote scientific research owing to high genetic similarity between primates and humans^22^. Although the potential importance and values of NHP models in basic research are indispensable, an organism-wide single-cell atlas is still pending for primates. Here we present a compendium of single-cell regulomic and transcriptomic data from *Macaca fascicularis* (cynomolgus monkeys) that comprises 40 distinct cell types from 16 organs and tissues, greatly extending our current view^23–25^ of single cell landscapes in this model species. This cell atlas -- which we denote ‘Monkey Atlas’ -- represents a new resource for NHP cell biology.

## Results

### Mapping a cynomolgus monkey multi-organ cell atlas by multi-omic analysis

To generate a reference cell map of monkey, we performed both single-cell RNA sequencing (scRNA-seq; 10x Genomics; n=174,233) and scATAC-seq (10x Genomics; n=66,566) on more than 240,000 high-quality cells from 16 tissues and organs in one male or/and one female cynomolgus monkeys (**Fig. 1a** and Supplementary Fig. 1). We integrated all of the scRNA-seq data using canonical correlation analysis (CCA)^26^ to correct for batch effects. Unsupervised clustering based on t-distribution stochastic neighbor embedding (t-SNE) resolved major cell types, including epithelial, ciliated epithelial, mesenchymal, immune, endothelial, spermatid, and secretory cell populations. These cells could be subdivided into 40 transcriptionally distinct clusters with cluster-specific markers (**Fig. 1b,c**). Due to technical and financial constraints, not every organ was analyzed in each monkey or by data modality. Nevertheless, the overall gene expression patterns or cell composition for the same organs or functional related organs (e.g., stomach, liver, spleen and colon from the digestive system) are quite similar (Supplementary Figs. 2 and 3); the analysis of multiple organs from the same monkey enable us to obtain data that is controlled for uncertain effects (such as age, sex, diet, environment and so on).

**Figure 1.**
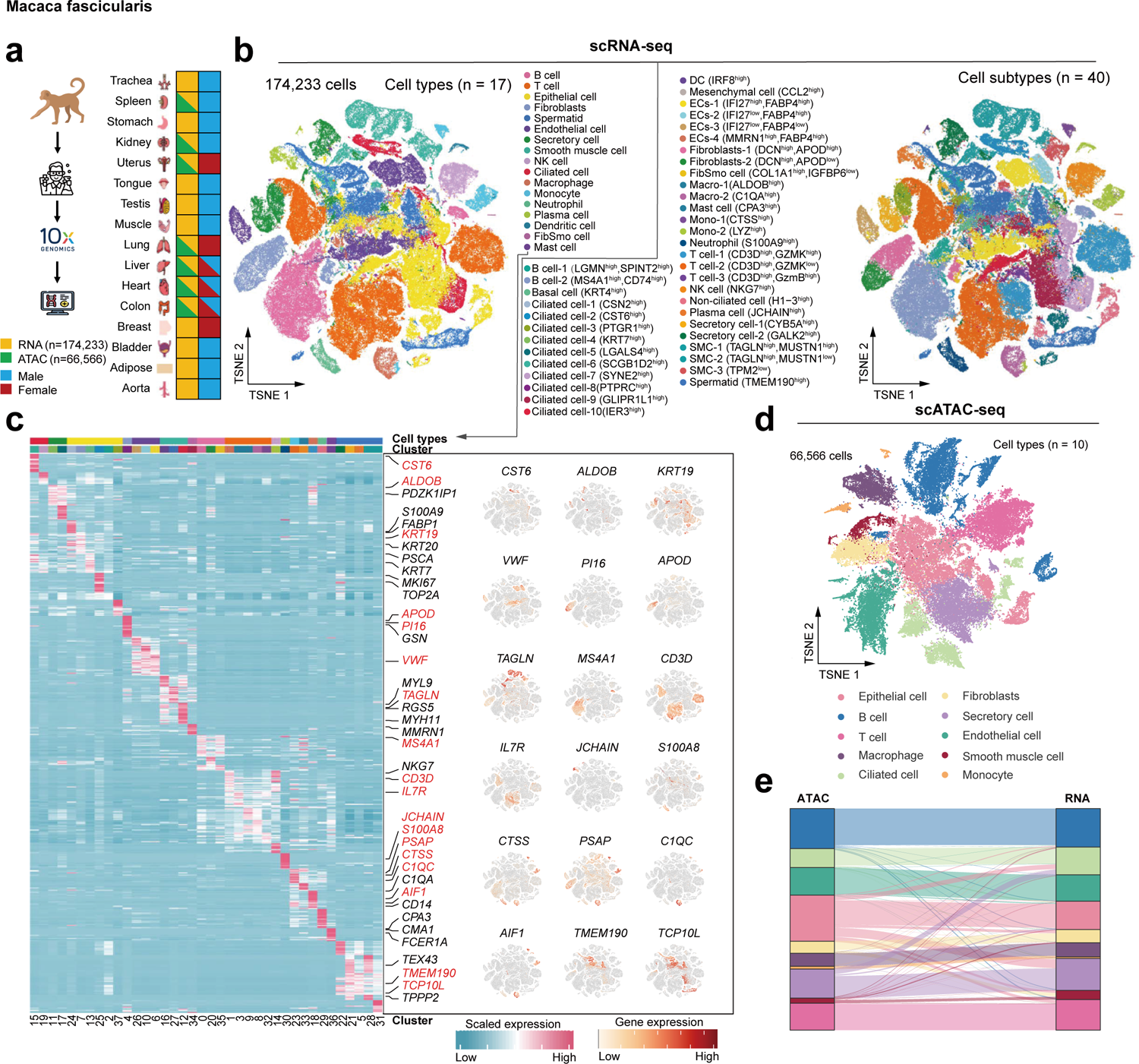
Single-cell landscapes of 16 organs from cynomolgus monkeys. **(a)** Workflow of sample collection and single-cell transcriptome analysis for 16 organs and chromatin accessibility analysis for 7 organs from normal cynomolgus monkey. **(b)** Cell type identification (scRNA-seq), including 174,233 cells, 17 major cell types, and 40 cell subtypes. TSNE of cells were either colored by major cell types (left) or colored by cell subtypes (right). **(c)** Heatmap with the scaled expression levels of cell type-specific marker genes (left). 18 marker genes expression were randomly selected to exemplify the specificity of marker gene in the right. **(d)** Overview of cell type identification in scATAC-seq analysis. 66,566 cells and 10 major cell types were identified and cells were colored by major cell types. **(e)** Sankey diagram of scRNA-seq and scATAC-seq data cell-type mapping.

To analyze scATAC-seq cells from the organs of liver, colon, uterus, spleen, lung, heart and kidney, we created a count matrix of fragments across the genome. We demonstrated the overall high quality of scATAC-seq data based on the enrichment analysis of accessible DNA sequences relative to the transcriptional start site (TSS) and the size distribution of unique fragments (Supplementary Fig. 4). T-SNE clustering analysis of scATAC-seq data revealed ten major cell types annotated based on chromatin accessibility at the promoter regions of well-characterized marker genes (**Fig. 1d**). For the organs with matched scRNA-seq and scATAC-seq data, we performed cross-modality integration analysis using mutual nearest neighbors (MNNs) approach (Supplementary Fig. 5; see **Methods**). We assigned cell type cluster labels from matched scRNAseq data to scATAC-seq cells. This revealed that cell types identified by scRNA-seq and scATAC-seq are highly consistent (**Fig. 1e**), highlighting the quality of the dataset.

### Epithelial cell heterogeneity and developmental dynamics across organs

Epithelial cells account for the largest part in the integrated cell map. To dissect epithelial heterogeneity, we extracted epithelial cells and performed unsupervised sub-clustering analysis (**Fig. 2a**). The analysis identified 14 cell clusters (E01-E14), including basal cells, secretory cells, ciliated cells and non-ciliated cells, according to distinct pattern of marker gene expression (**Fig. 2a,b**). It is worth noting that we observed a large proportion of ciliated epithelial cells in various tissues (**Fig. 2c** and Supplementary Fig. 6a,b). Ciliated cells play an important role in cleaning pathogenic microorganisms and signal transduction^27^. Gene ontology (GO) enrichment analysis based on differentially expressed genes revealed ciliated cell subpopulations with different biological functions. For example, subpopulations of E09 (*SCGB1D2*^high^ ciliated cells) and E12 (*GLIPR1L1*^low^) enriched for pathways related to cellular response to stimulus, cell communication and intracellular signal transduction, whereas E05 subpopulation related to RNA biosynthetic process (**Fig. 2d**).

**Figure 2.**
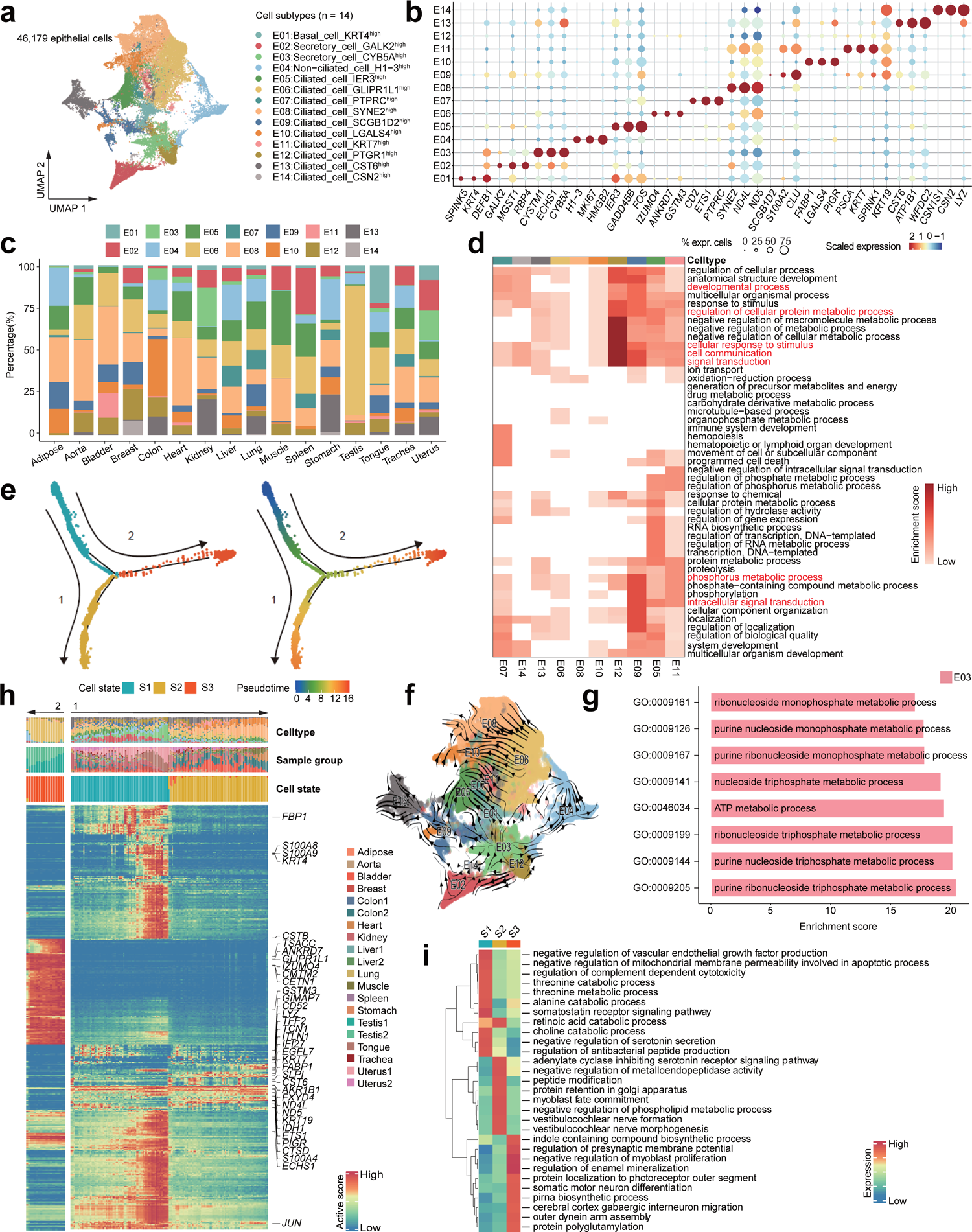
The heterogeneity and developmental state of epithelial cells. **(a)** Distribution of 14 epithelial cell subtypes on the UMAP. **(b)** Dot plot shows representative differentially expressed genes (DEGs) across epithelial subtypes. The size of dot is proportional to the fraction of cells which express specific genes. Color intensity corresponds to the relative expression of specific genes. **(c)** Bar plot shows the percentage of cell subtypes in each organ. **(d)** Heatmap shows functional pathways of ciliated epithelial cell enrichment, darker color represents a higher enrichment score. **(e)** Semi-supervised pseudotime trajectory of subtypes (E01-E14) of epithelial cells by monocle2. Trajectory is colored by cell states (left) and pseudotime (right). **(f)** Unsupervised pseudotime trajectory of subtypes (E01-E14) of epithelial cells by RNA velocity. Trajectory is colored by cell subtypes. Arrowhead direction represents the trend of cell pseudo-temporal differentiation. **(g)** Bar plot shows functional pathways of E03 subtype. **(h)** Heatmap showing the scaled expression of differentially expressed genes across pseudotime from (**e**). Genes (on the right of the heatmap) are assigned to specific cell states based on their expression levels. Bar plots at the top of the heatmap are scale diagrams of different cell types, samples and cell states during pseudotime differentiation trajectory. **(i)** Heatmap showing functional pathways enriched in three cell states (S1, S2, S3) by GSVA analysis.

To explore the developmental dynamics of epithelial cells, we performed pseudotime trajectory analysis using both Monocle2^28^ and RNA velocity^29^. We determined the cluster E03 (*CYB5A*^high^ secretory cells) as the start point of trajectory based on estimated latent time by RNA velocity (**Fig. 2f**). Accordingly, highly expressed genes in E03 are functional related to ATP metabolic process and purine ribonucleoside monophosphate metabolic process (**Fig. 2g**). Epithelial cells were arranged into a trajectory with two bifurcations and three cell states with E03 as the root (**Fig. 2e**). It is worth noticing that ciliated epithelial cells are in different states of differentiation. For example, cluster E06 was predominant in bifurcation 2 (**Fig. 2h** and Supplementary Fig. 6e). Some marker genes are not in the E06 subgroup, such as *CST6*, *PDZK1IP1*, *KRT19*, and *PSCA*, they had low relative expression in state 3 (Supplementary Fig. 6d). From the perspective of sample type, almost all of the epithelial cells of testis tissue are present in bifurcation 2 (major in state3) (**Fig. 2h** and Supplementary Fig. 6c). Differential genes such as *GABRR1*, *GRIK2*, *CEP41*, *CFAP20*, *TPGS1* in state3 showed enrichment in presynaptic membrane potential and protein polyglutamylation (**Fig. 2i**). This may be due to the ability of testis to produce sperm and male hormones. We speculate that GLIPR1L1^high^ ciliated cells are the main effector epithelial cells of the testis.

### Stromal cellular heterogeneity

Stromal cells are an important component of body tissues^30^. In the stromal compartment, we identified 11 clusters (S01-S11) belonging to four major cell types including endothelial cells, fibroblasts, FibSmo cells and smooth muscle cells (**Fig. 3a-c** and Supplementary Fig. 7a). Although these cell clusters were identified in all organ tissues, the heterogeneity of stromal cells was observed in different organs (**Fig. 3d,e** and Supplementary Fig. 7b). Most mesenchymal cells were generated from kidney; almost all fibroblasts in testis are from S05 (DCN^high^APOD^high^ fibroblasts); there are a large number of TAGLN^high^MUSTN1^low^ smooth muscle cells in aorta tissues but few other smooth muscle cells (**Fig. 3d**). Considering that stromal cells have a certain differentiation potential^31^, we applied RNA velocity analysis to explore developing states of stromal cells (**Fig. 3f**). The results show that fibroblasts have the capacity of differentiation to smooth muscle cells and endothelial cells (**Fig. 3g,i** and Supplementary Fig. 7c-e). It is worth noting that fibroblasts of cynomolgus monkeys have a strong metabolic ability rather than the ability to synthesize collagen (**Fig. 3h**).

**Figure 3.**
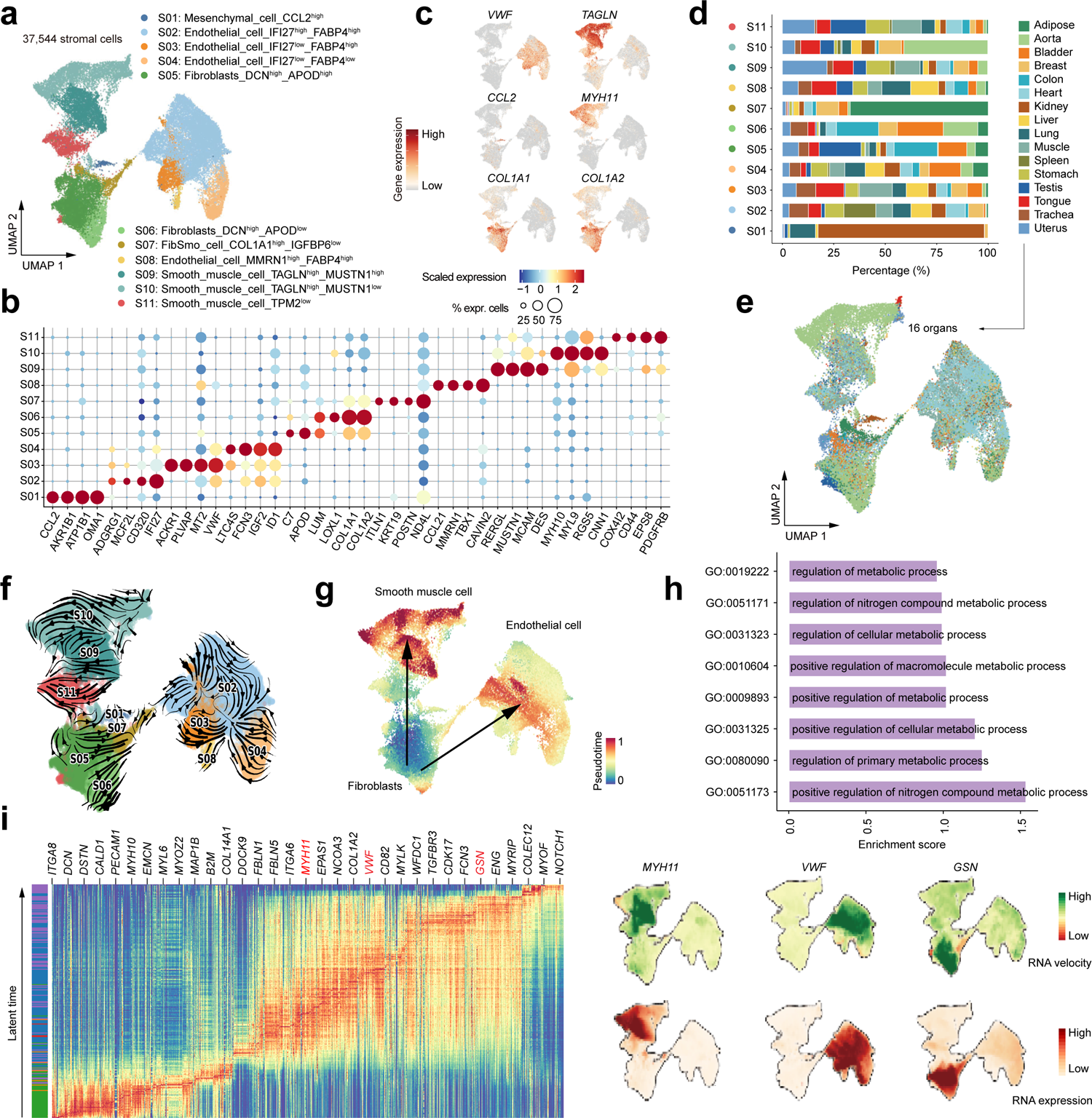
The heterogeneity of stromal cells. **(a)** Distribution of 11 stromal cell subtypes on the UMAP. **(b)** Dot plots showing representative top differentially expressed genes (DEGs) across stromal subtypes. Dot size is proportional to the fraction of cells expressing specific genes. Color intensity corresponds to the relative expression of specific genes. **(c)** Feature plot shows the expression of marker genes. **(d)** Bar plot showing the percentage of cell subtypes in each organ. **(e)** Distribution of stromal cells in different organs on the UMAP. **(f)** Unsupervised pseudotime trajectory of subtypes (S01-S11) of stromal cells by RNA velocity. Trajectory is colored by cell subtypes. The arrow direction is the trend of cell pseudo-temporal differentiation. **(g)** The UMAPs showing the pseudotime differentiation trajectory of fibroblasts, smooth muscle cells and endothelial cells. **(h)** Bar plot shows functional pathways of fibroblasts. **(i)** Heatmap showing the scaled expression levels of cell cell type-specific marker genes along pseudotime differentiation trajectory. Examples of marker expression are shown in the right UMAPs.

### Heterogeneity of immune cells

Immune cells are essential for maintaining body homeostasis^32^. We identified 72,284 immune-related cells from the investigated organs, including B cells, T cells and myeloid cells, and these cells were further grouped into 13 major clusters (I01-I13) based on known or novel gene signatures (**Fig. 4a,b** and Supplementary Fig. 8a,b). Although the annotated immune cell clusters can be found in all organs (**Fig. 4c,d**), the relative proportion of immune cells varied greatly in different organs. For example, We noticed that NKT_cell_CD3D^high^_GZMK^high^_GzmB^high^ cells vary widely in muscle tissues compared to other tissues (**Fig. 4e**). Subsequently, we analyzed differentially expressed genes of NKT_cell_CD3D^high^_GZMK^high^_GzmB^high^ cells in muscle tissues. We observed that mitochondria-related genes (ATP6, COX3 and ND1) were the main differentially expressed genes in NKT_cell_CD3D^high^_GZMK^high^_GzmB^high^ cells of muscle tissue compared to other tissues (**Fig. 4f** and Supplementary Fig. 8c).

**Figure 4.**
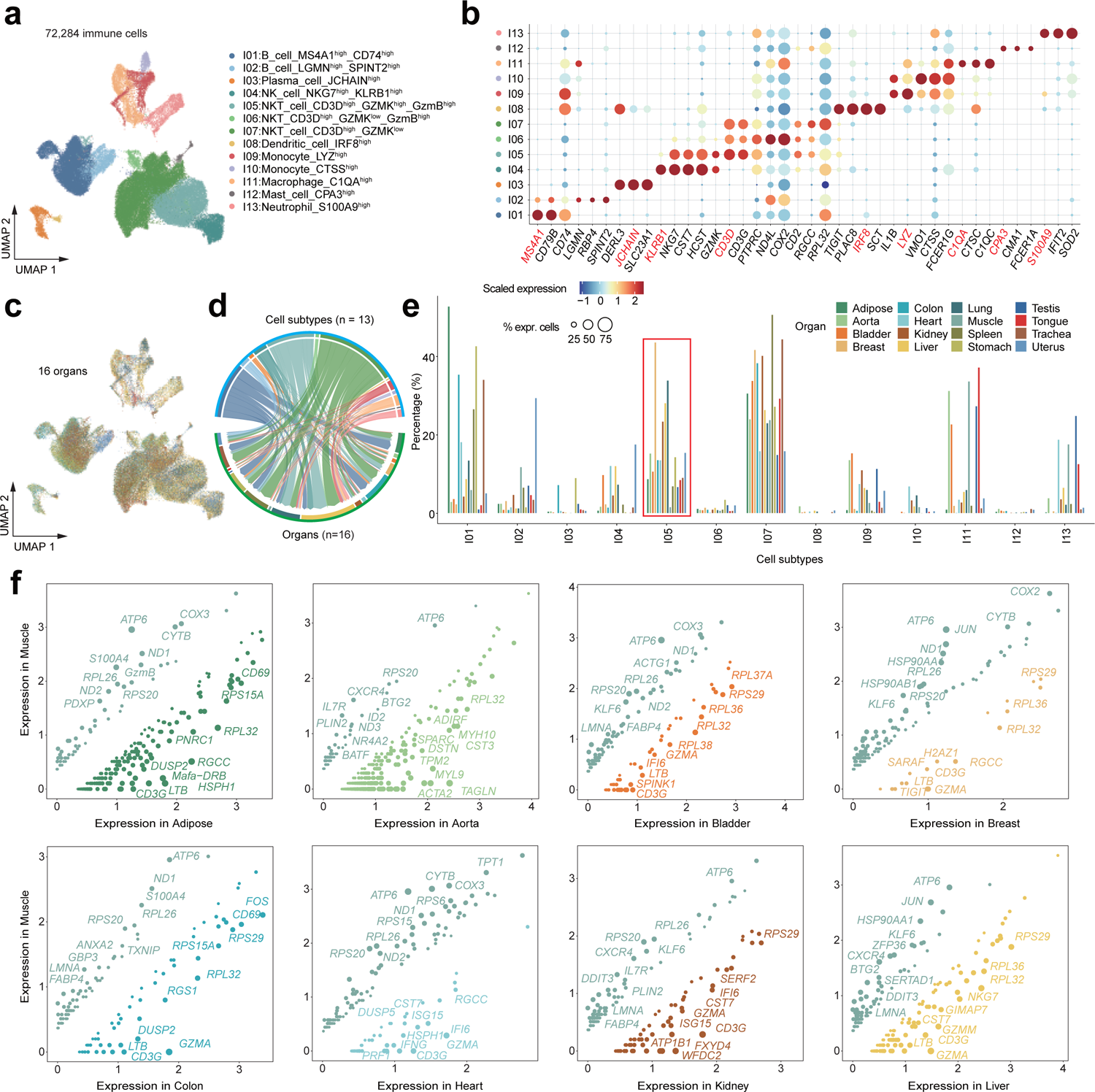
Immune cell heterogeneity. **(a)** Distribution of 13 major immune cell types on the UMAP. **(b)** Dot plots shows representative top differentially expressed genes (DEGs) across stromal subtypes. Dot size varies synchronously with the fraction of cells expressing specific genes. Color intensity corresponds to the relative expression of specific genes. **(c)** Distribution of immune cells in different organs on the UMAP. **(d)** Chord diagram maps the different cell types in all organs as a whole. The width of the arrow represents the proportion of cell types. **(e)** The bar chart shows the proportion of different cell types in the same organ. **(f)** Scatter plots shows pairings of gene expression between muscle and other organs. Each point represents a DEG, and its size is proportionate to the fold change.

### Dynamics of cell-cell interactome

To decipher the dynamics of intercellular communications in different tissues, we employed CellPhoneDB to identify potential ligand-receptor pairs among the major cell types. We observed that there are strong intercellular interactions among stromal cells, epithelial cells and myeloid cells (**Fig. 5a,b**). Generally, the intensity and pattern of cellular interactions between cells are tissue-specific (**Fig. 5c**). For example, tongue and uterus tissues show stronger cellular interactions, while intercellular interactions in testicular tissue is weaker than other tissues (Supplementary Figs. 9 and 10). To chart the rewiring of molecular interactions regulating cell-cell interactions, we mapped ligand-receptor pairs in specified cell subpopulations in different organs (**Fig. 5d**). In brief, the “CD99-PILRA” ligand-receptor pair is specific in the interaction between stromal cells and myeloid cells, particularly in adipose, aorta, and colon. As an inhibitory receptor of immunoglobulin-like type 2 receptor (PILR), PILRA has been shown to bind to the CD99 ligand for immune regulation^33^. The “CCL4L2-VSIR” pair occurred exclusively in the interaction of myeloid cells and T cells. In contrast, the “LGALS9-CD44” pair contributed to most immune cell related interactions. Accordingly, CD44 plays a role in innate immunity and subsequent adaptive responses, and has extensive inflammatory and proliferative effects on a variety of cell types^34, 35^. Taken together, these results predicted the possible molecular mechanisms underlying cell-cell communication in various tissues (**Fig. 5d**).

**Figure 5.**
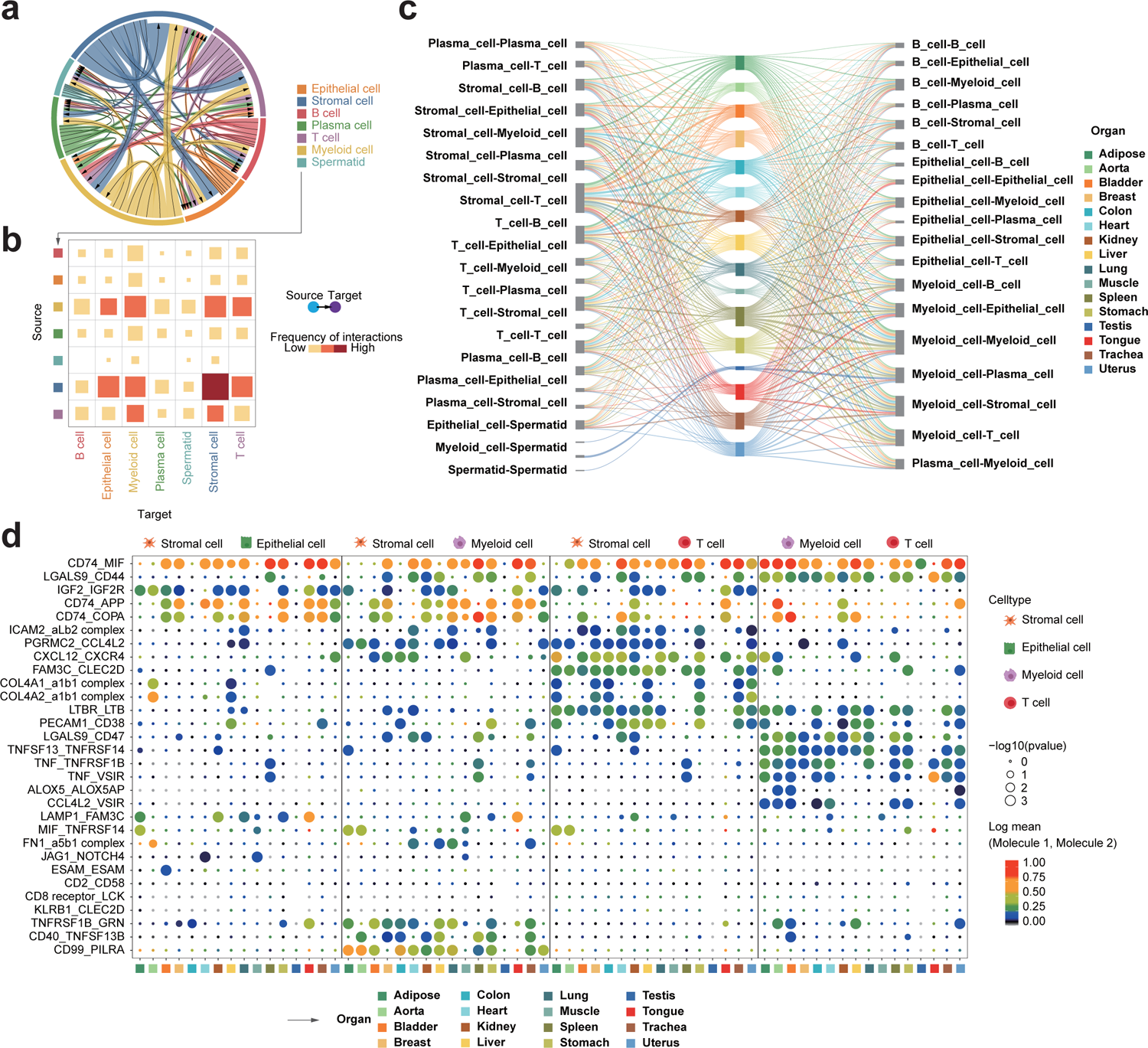
Dynamics of cell-cell communication networks. **(a)** Chordal diagram of the integrated cell-cell interaction network among the major cell types of 16 organs. **(b)** Heatmap shows interaction intensity of cellular interactome from (a). Block sizes and colors are proportional to the interaction frequency. **(c)** Sankey diagram shows the cell-cell interactions of different cell types in 16 organs. The thickness of lines represents the strength of cell-cell interactions. **(d)** Dot plot of interactions between selected cell subtypes in different organs. Each row represents a ligand-receptor pair, and each column defines a cell-cell interaction pair.

### Single-cell chromatin landscape of major organs in cynomolgus monkey

To deconstruct the gene regulation principles of complex tissues in cynomolgus monkey, we examined the single-cell chromatin accessibility landscape of major organs including colon, kidney, lung, uterus, heart, liver and spleen by scATAC-seq. In total, we generated scATAC-seq profiles from 66,566 cells after quality control. We identified 22 distinct cell clusters in the integrated cell map according to cluster-specific *cis*-elements and visualized single-cell profiles with uniform manifold approximation and projection (UMAP) (**Fig. 6a** and Supplementary Fig. 11a). For example, clusters 1-4 demonstrated accessibility at *cis*-elements neighboring B cell genes, including *CD22*, *MS4A1* and *TNFRSF13C*, while the cluster 22 demonstrated accessibility at *cis*-elements neighboring T cell genes, including *CD3D* and *IL7R* (Supplementary Fig. 12). We detected 397,773 *cis*-elements across all clusters, ranging from 3,046 to 75,001 peaks in each clusters (**Fig. 6b**). As expected, most of *cis*-elements were derived from promoters, intronic or distal intergenic regulatory regions. We observed that most cell clusters are organ-specific (**Fig. 6c**) and cluster-specific *cis*-elements exhibited organ-specific accessibility accordingly (**Fig. 6d**).

**Figure 6.**
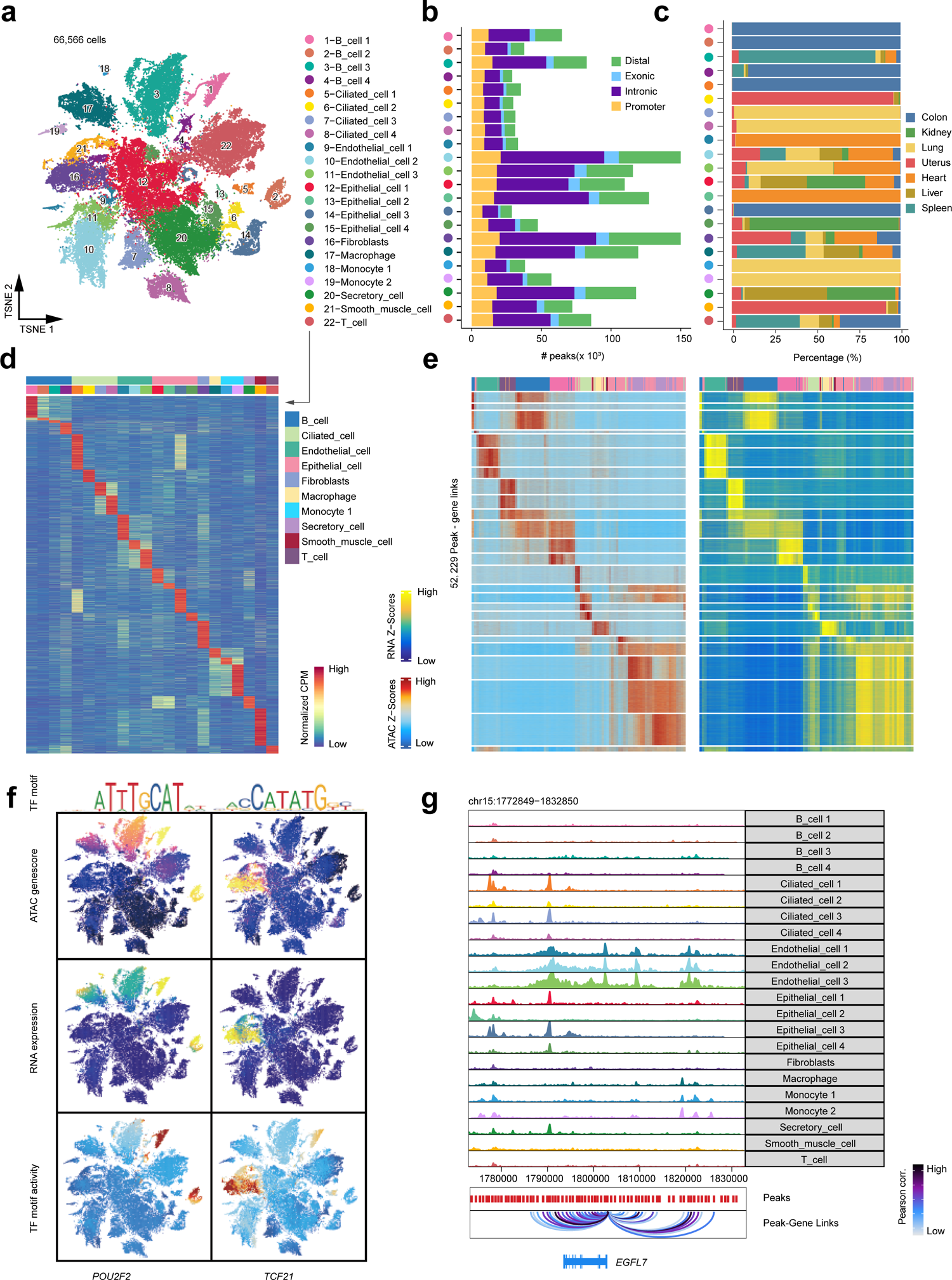
Single-cell chromatin landscape of major organs of cynomolgus monkeys. **(a)** Cell type identification (scATAC-seq), including 66,566 cells, and 22 cell subtypes. Shown is the tSNE of cells colored by cell subtypes. **(b)** Bar plot shows the number of reproducible peaks identified from each cluster. The peaks are classified to four category: distal, exonic, intronic and promoter. **(c)** Percentage of cell subtypes in each organ shown by bar plot. **(d)** Heatmap of 80,270 marker peaks across 22 subclusters identified by bias-matched differential testing (FDR <= 0.01 and Log2FC >= 3). **(e)** Chromatin accessibility and gene expression of 52,229 significantly (R >0.45 and FDR < 0.1) linked peak-gene pairs illustrated by heatmap. **(f)** Profile of POU2F2 and TCF21 gene accessibility, gene expression (inferred from scRNA-seq) and TF motif activity. **(g)** Visualization of the EGFL7 locus with the maximum number of peak-gene pairs shown by genome browser track (chr15: 1,772,849−1,832,850).

Comparison analysis of scATAC-seq and scRNA-seq data highlighted concordant patterns of chromatin accessibility and gene expression across clusters, exemplified by marker genes (*POU2F2* and *TCF21*) in specific cell types (**Fig. 6e,f**). We also computed TF deviation scores using chromVAR^36^, which measured the accessibility of TF binding “footprint” genome-wide in each single cell. Indeed, TF deviation scores for *POU2F2*, a B-cell-specific transcription factor involves in cell immune response by regulating B cell proliferation and differentiation^37, 38^, were increased in B cell clusters (**Fig. 6f** and Supplementary Fig. 11b). Similarly, the TF deviation scores for *TCF21*, an essential regulator of fibroblasts in development^39^, were increased in the fibroblasts cell cluster (**Fig. 6f** and Supplementary Fig. 11b). Furthermore, we applied Cicero^40^ to identify co-accessible *cis*-elements at genome-wide, as exemplified at the gene locus of *EGFL7* (**Fig. 6g**), an endothelium-specific secreted factor mostly produced by blood vessel endothelial cells during development^41–43^. We observed increased enhancer-promoter connections in endothelial cell clusters at the promoter of *EGFL7*. Overall, our scATAC-seq data provide a rich resource for unbiased discovery of cell types and regulatory DNA elements in cynomolgus monkey.

### Cell-type specific and organ-specific transcriptional gene regulatory networks

TFs are important regulators controlling cell identity and tissue-specific gene expression. To infer cell-type and organ-specific transcriptional regulatory programs based on the monkey cell transcriptional landscape, we applied SCENIC (single-cell regulatory network inference and clustering) to identify TF regulons. We identified several TF regulon modules that were active in either cell-type (n=8; Supplementary Fig. 13a,b) or organ-specific manners (n=6; **Fig. 7a,b**). Subsequently, we analyzed representative TF regulons across different cell types (n=7; **Supplementary Figs. 13c** and 15) or different organs (n=16; **Fig. 7c** and Supplementary Fig. 14). The identified TF regulons are highly cell-type or organ-specific based on regulon activity scores (**Fig. 7d** and Supplementary Fig. 13d). Finally, The representative TF regulons and their associated target genes were organized into cell-type specific or organ-specific gene regulatory networks (**Fig. 7e** and Supplementary Fig. 13e).

**Figure 7.**
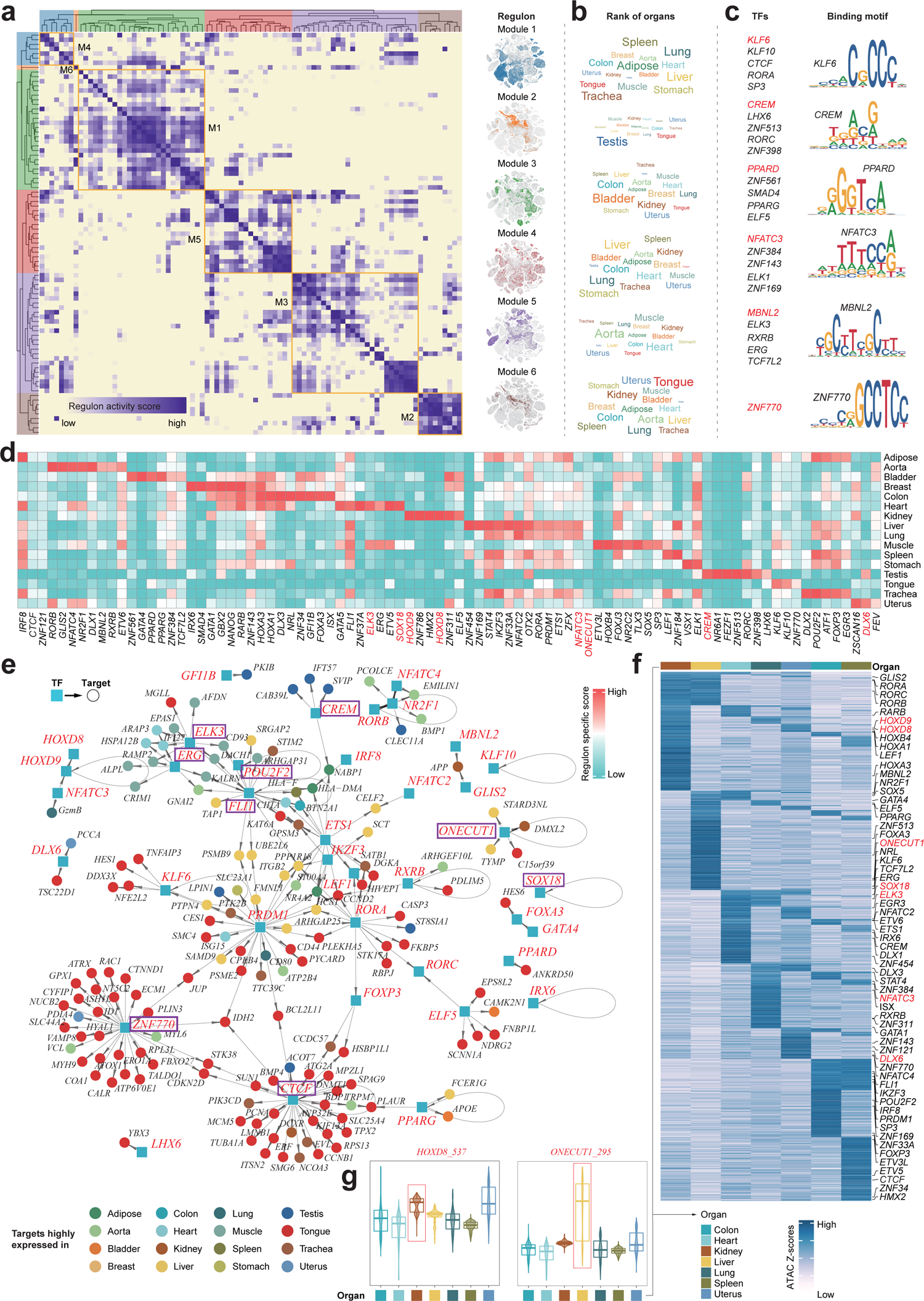
Organ-specific transcriptional regulatory networks. **(a)** Identification of regulon modules using SCENIC. Heatmap (left) shows the similarity of different regulons (n=87) based on the AUCell score. Eight regulon modules were identified based on regulon similarity. UMAPs (right) illustrate the average AUCell score distribution for different regulon modules (in different colors). **(b)** Wordcloud plots shows enrichment of organs in different regulon modules. **(c)** Representative transcription factors and corresponding binding motifs in different regulon modules. **(d)** Heatmap showing transcription factors enriched in different organs. Color depth represents the level of regulon-specific score. **(e)** Integrated gene-regulatory networks of the regulons. Regulon-associated TFs are highlighted in blue rectangles and target genes in circles. Target genes (in circles) are colored according to their highly expressed organs. **(f)** Heatmap showing gene-activity scores of marker genes in the indicated organs. **(g)** Violin plot showing motif activity (measured by TF chromVAR deviations) of TF regulons highlighted in (**f**).

In the cell-type specific gene regulatory networks, we observed that CREM specifically controls proliferation-related target genes such as DAZL and HEY2 in spermatid cells. Immune-related TFs such as IRF2, FLI1 and IK2F3 are shown to regulate immune cell identity genes such as S100A4 and CD48. FEV, a known TF that regulates the development of hematopoietic stem cells^44^, extensively link to target genes actively expressed in immune cells and epithelial cells (Supplementary Fig. 13e).

In the organ-specific gene regulatory networks, we found that target genes of ZNF770 and CTCF are specifically expressed in tongue. The spermatid cell-specific TF regulon CREM regulated genes actively expressed in testis. ETS-factors (ELK3, ERG, and FLI1) together with pre-/immature-B TFs (POU2F2) positively regulated genes showed elevated activity in heart and muscle (**Fig. 7e**).

To further confirm the unbiased inference of organ-specific TF regulons based on scRNA-seq data (see **Fig. 7d**), we validated the organ-specific TF regulons using organ-matched scATAC-seq data. We therefore measured chromatin accessibility at *cis*-elements containing a specific TF binding motif using chromVAR^36^ and accessibility changes were analyzed in binding sites for the above identified organ-specific TFs (denoted as TF deviation scores). In general, TF deviation scores showed similar organ-specific patterns to regulon activity scores (**Fig. 7f**). For example, the HOXD8 regulon is kidney-specific and showed high TF deviation scores in kidney, while the regulon activity and TF deviation scores of ONECUT1 are both liver-specific (**Fig. 7g**). These analyses emphasize the unbiased prediction of organ-specific gene regulatory networks at the single-cell level.

### Comparison of cell landscapes among human, mouse and cynomolgus monkey

The cynomolgus monkey cell landscape offers the opportunity to compare the cellular components and transcriptomic dynamics across species with similar organ compositions. Here we integrated scRNA-seq data from the non-human primate cynomolgus monkey (by this study), human^16^ and mouse^45^ with matched organs/tissues using orthologous genes for cross-species analysis (**Fig. 8a** and Supplementary Fig. 16a,b). The integrated cell map consists of 338,932 cells (**Fig. 8b**), which were grouped into 15 major cell types (**Fig. 8c** and Supplementary Fig. 16c,d). Although the cell-type compositions largely varied in the three species (**Fig. 8d,i**), the expression patterns of representative marker genes and transcriptomic similarity of cell types were overall consistent across species (**Fig. 8e,f** and Supplementary Fig. 14e,f). Consistent with previous single-cell comparative analyses^3, 18–20^, the gene expression patterns of the major cell types are conserved in all three species, including immune, stromal and epithelial cells (**Fig. 8g**). As expected, cynomolgus monkey and human showed significantly higher cell-type similarity in orthologous gene expression than comparisons in other species (**Fig. 8h**).

**Figure 8.**
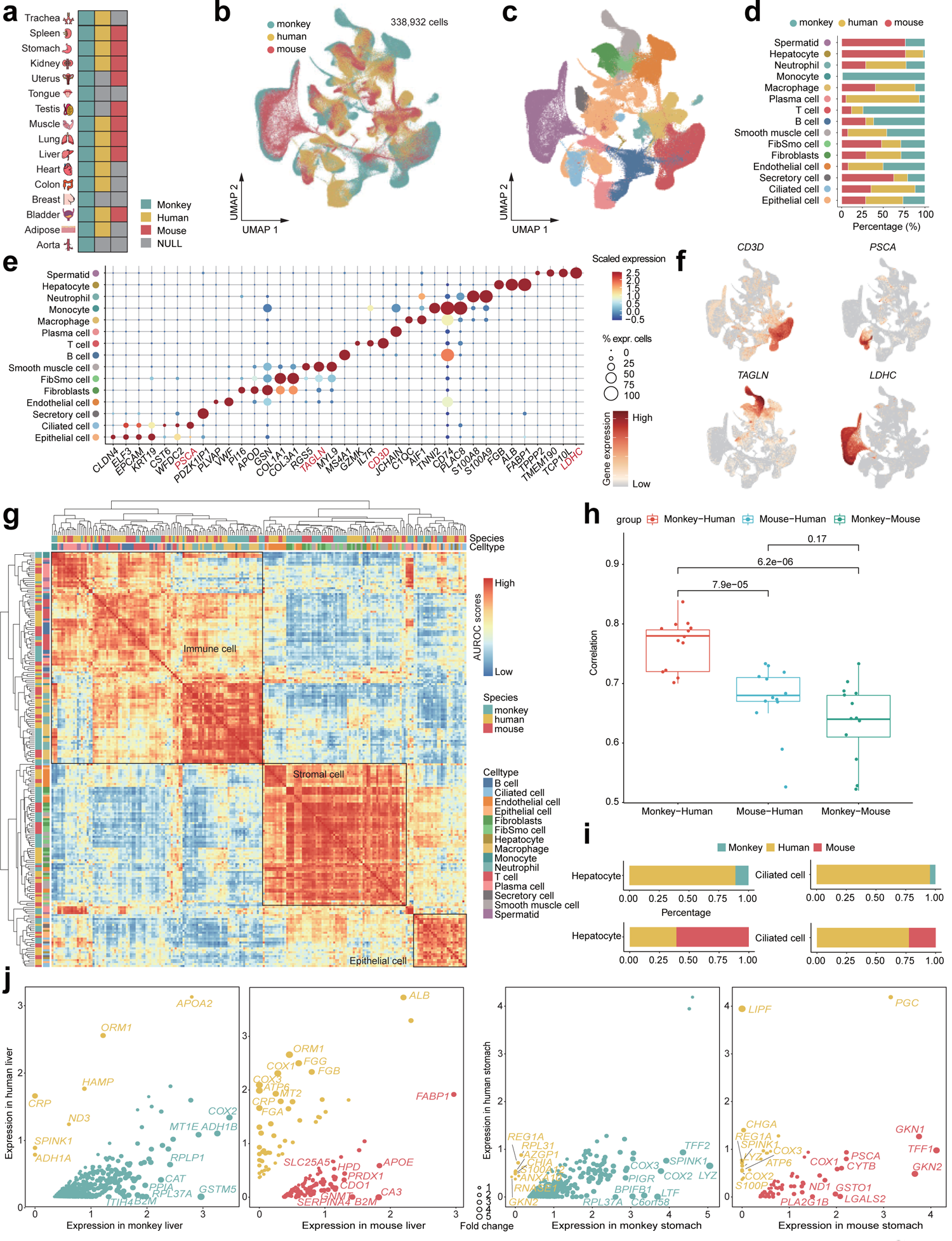
Comparison of cell landscapes of human, mouse and cynomolgus monkey. **(a)** Integration of data from 16 organs from cynomolgus monkey, 11 organs from human, and 9 organs from mouse. **(b)** Distribution of cells from cynomolgus monkey, human and mouse on the UMAP. **(c)** Distribution of 15 major cell types on the UMAP. **(d)** Bar plot showing the percentage of cell types in cynomolgus monkey, human and mouse. **(e)** Dot plots showing representative top differentially expressed genes (DEGs) across cell types. Dot size is proportional to the fraction of cells expressing specific genes. Color intensity corresponds to the relative expression of specific genes. **(f)** Feature Plot showing the expression of marker genes. **(g)** Correlation of orthologous gene expression between human, mouse and monkey pseudo-cell types (n=212) based on AUROC scores. The AUROC scores are calculated by MetaNeighbor to measure the similarity of cell type. The hierarchical clustering is calculated with pseudo-cell types. **(h)** Box plot showing correlation of top genes expression in cynomolgus monkey, human and mouse integrated data (Spearman method). Dots represent different cell types. **(i)** The proportion of hepatocytes in liver and ciliated cells in stomach of cynomolgus monkey, human and mouse samples. **(j)** Scatter plots showing a pairwise comparison of gene expression across species organs. Each point represents a DEG, and its size is proportionate to the fold change. Two scatter dots on the left show DEGs of Hepatocyte in the cynomolgus monkey, human and mouse samples, and Ciliated cells on the right.

We next investigated the transcriptomic dynamics of the same cell types among different species, with a specifical focus on varying cell types across species such as hepatocytes and ciliated cells (**Fig. 8i**). To this end, we performed pairwise comparison of gene expression in liver for hepatocytes and in stomach for ciliated cells (**Fig. 8j**). In the differential analysis of gastric ciliary cells between human and cynomolgus monkey, we found that *LYZ* was highly expressed in cynomolgus monkeys^46^. *LYZ* has a dual role of immune defense and digestive function^47^. As a gastric lipase, *LIPF* is expressed in human chief cells and promotes lipid metabolism^48^. *ORM1*, as an acute-phase protein, was highly specifically expressed in human hepatocytes and had a certain promoting effect on liver regeneration.

## Discussion

Non-human primates (NHP) are similar to humans in terms of anatomy, physiology and biochemical metabolism. Cynomolgus monkeys, a well-established laboratory animal model, have outstanding contributions to the scientific field^49^. Although several single cell transcriptomic atlases have recently been established in cynomolgus monkeys based on a few organs (including ovary, lung, heart and artery) ^23–25^, an organism-wide single-cell map is still lacking in this model species. Here, we chart a reference cell map of cynomolgus monkeys (named ‘Monkey Atlas’) using both scATAC-seq and scRNA-seq data across multiple organs, allowing us to gain deeper insights into the molecular dynamics and cellular heterogeneity of the cynomolgus monkeys organism.

As a proof of concept, we have performed various analyses based on the Monkey Atlas to show its wide uses, including the discovery of new putative cell types, the identification of key regulators in organ specification, and the ability to compare cell types across organs and species. For instance, our data shows that ciliated cells present in various organs of cynomolgus monkeys and the different ciliated cell subpopulations show various functions related to metabolic process, signal transduction, and cellular response to stimulus (**Fig. 2**). This observation somehow expands our notion that ciliated cells are generally found in respiratory system^50^ with vital role in cleaning pathogenic microorganisms and signal transduction^27^.

Recently, comprehensive reference cell maps across organs have been established in human^3, 51^ and mouse^45, 52^. Although the Monkey Atlas does not provide exhaustive characterization of all organs in cynomolgus monkeys, it does offer a rich dataset of the most populously studied organs in biology. In this regard, we performed cross-species integration analysis of cell maps to explore the molecular and cellular differences among the three species with comprehensive single cell data. We noticed that cynomolgus monkeys and human both have abundant immune cells and epithelial cells and a comparative composition of cell types in matched organs. This indicates that cynomolgus monkeys are ideal models to study complex diseases.

In conclusion, the Monkey Atlas provides valuable information about the most populous and important cell populations in NHP, and presents a foundation for preclinical studies.

## Methods

### Organ tissue collection

The cynomolgus monkey sample collection and research conducted in this study were approved by the Research Ethics Committee of the Changchun Biotechnology Development Co., Ltd. (Approval Number: 21001). Tissues were collected from 16 organs including trachea, spleen, stomach, kidney, uterus, tongue, testis, muscle, lung, liver, heart, colon, breast, bladder, adipose and aorta. To be more specific, tissues were cut into 1-2 mm^3^ pieces in RPMI-1640 medium (Gibico) with 12% fetal bovine serum (FBS, Gibico). Then the tissues were enzymatically digested with gentleMACS (Miltenyi) according to manufacturer’s instruction. Cells were passed through a 70 μm cell trainer (Miltenyi) and centrifuged at 300 g for 5 min at 4 ℃. The pelleted cells were re-suspended in red blood cell lysisbuffer (Beyotime) and incubated 1 min to lyse red blood cells. After wash twice with 1XPBS (Gibico), the cell pellets were re-suspended in sorting buffer (PBS with supplemented with 1% FBS). The single cells were captured in the 10× Genomics Chromium Single Cell 3’ Solution, and RNA-seq libraries were prepared following the manufacturer’s protocol (10× Genomics). The libraries were subjected to high-throughput sequencing on the Novaseq6000 platform, and 150-bp paired-end reads were generated.

### scRNA-seq and data processing

Single-cell gene expression data were aligned to the *Macaca fascicularis* reference genome (macFas6) and processed for barcode assignment and unique molecular identifier (UMI) counting using the CellRanger v3.1.0 pipeline (10x Genomics). Filtered count matrices from the CellRanger pipeline were converted to sparse matrices using Seurat package (v4.0.0) in R^53^, and cells expressing more than 4000 genes or less than 200 genes and more than 20% of mitochondrial genes expressing in UMI counts were filtered out before downstream analysis. Filtered data were then log normalized and scaled, with cell-cell variation due to UMI counts and percent mitochondrial reads regressed out. Then, we log normalised and scaled the filtered data to avoid cell-to-cell variation caused by UMI counts and percent mitochondrial reads removal.

As the samples involved the integration of large multi-organ samples such as trachea and spleen, Seurat’s Robust Principal Component Analysis (RPCA) method was adopted for data integration. Cell clustering was performed at 0.8 resolution using the “FindClusters” function, and cell identity were defined using the top 20 principal components (PCs), and 17 clusters were identified. Dimensionality was reduced by the “RunUMAP” function and by visual Uniform Manifold Approximation and Projection (UMAP). Different types of cells were extracted for subgroup cell clustering, and their first 20 PCS were used for clustering. In the end, we identified 40 different subgroups. To ensure the accuracy of subsequent analysis, all 40 different subgroups were processed to remove double cells. Wilcoxon rank-sum test (FindAllMarkers function with default parameters) was used to identify markers for each cluster. Marker genes for each cluster are shown in Supplementary Table S1.

### Creating a reference package for Macaca fascicularis

First, the FASTA and GTF files of the *Macaca fascicularis* reference genome were downloaded from the Ensembl database (version 6.0). Then, GTF files were filteded since it contains entries for non-polyA transcripts that overlap with protein-coding gene models. Because of the overlapping annotations, these entries can cause reads to be flagged as mapped to multiple genes (multi-mapped). Therefore, these entries were removed from the GTF file. Finally, we create a Reference Package using genome FASTA and filtered GTF files.

### Gene-set enrichment analysis

Gene-set enrichment analysis (GSEA) is a gene-based enrichment analysis method. The clusterProfiler package^54^ in R was performed in all the gene-set enrichment analyses in this study.

### Gene-set variation analysis

Gene-set variation analysis (GSVA)^55^ starts from gene expression amount and multiple pathway information. Unsupervised samples were classified according to pathway activity changes. Three states defined by pseudo time analysis were choose as groups and genes corresponding to the three states were utilized to conduct Gene-set variation analysis in R.

### Trajectory analysis using Monocle v.2

R package Monocle2 (version 2.99.3) was used to illustrate the epithelial cells state transition. In general, UMI count matrix of epithelial cells, cell phenotype information and gene annotation information, and the negbinomial.size() parameter were used to create a CellDataSet object. The variable genes obtained from epithelial cell types were detected by Seurat to sort cells in pseudotime. We used the DDRTree method and orderCells function for dimensional reduction and cell ordering. The secretory_cell_CYB5Ahigh cluster (E03) was defined as the root state argument and aligned via the “orderCells” function.

### Trajectory analysis using RNA velocity

We sorted the possorted_genome_bam.bam file in the outs folder generated after each organ sample ran CellRanger, and then use Velocyto’s run10x function to generate loom files. To run run10x, we needed to prepare rmsk.gtf and the genes.gtf of the genome as input files. In this paper, RNA Velocity was used for trajectory analysis of epithelial cells and stromal cells.

### Cell-cell interaction analysis

Cell-cell interactions among the cell types were estimated by CellPhoneDB (v2.1.1)^56^ with default parameters (10% of cells expressing the ligand/receptor) and using version 2.0.0 of the database. We converted the normalized *Macaca fascicularis* genes into human homologous and used them as inputs. Interactions with p-value < 0.05 were considered significant. We considered only ligand-receptor interactions based on the annotation from the database, for which only and at least one partner of the interacting pair was a receptor, thus discarding receptor-receptor and other interactions without a clear receptor.

### Create cisTarget databases for Macaca fascicularis

Firstly, chromosomes were cut at start and stop sites. EnsemblID, SymbolID, and annotation information from the GTF file were extracted to make the genes.bed file. The fasta file was generated from genes.bed file (10kb up- and 10kb downstream of the TSS) with ‘bedtools getfasta’. Human motifs were downloaded from the CIS-BP database. Then, we extracted Motif_ID and TF_Name from the motifs file and outputted them as motifs_list. After, we converted all motif files in the pwms_all_motifs folder into motif.cb files. Finally, the fasta file, motifs_list file, and motif.cb files were used as input files to create cisTarget databases for Macaca fascicularis(macFas6).

### Gene regulatory network

To identify cell type-specific gene regulatory networks, we performed Single-cell Regulatory Network Inference and Clustering (v0.10.0; a Python implementation of SCENIC)^3^ in our *Macaca fascicularis* dataset. First, the original expression data were normalized by dividing the gene count for each cell by the total number of cells in that cell and multiplying by 10,000, followed by a log1p transformation. Next, we used the normalized counts to generate the co-expression module, using the GRNboost2 algorithm implemented in the arboreto package(v.0.1.3). Finally, we used pySCENIC with its default parameters to infer co-expression modules, cis-regulatory were used to filter motif analysis (RcisTarget, using macFas6 motif set), in order to only retain the corresponding transcriptional speculative direct binding target module enrichment factor (TF). The rest of the modules were trimmed due to the lack of motif target support. An AUCell value matrix was generated to represent the regulators in each cell with different activity. The final gene regulatory networks consisted of 86 regulons for our *Macaca fascicularis* dataset as shown in figure 7a. GRNs can be visualized using igraph package in R.

### scATAC-seq data pre-processing

The scATAC-seq sequencing data are pre-processed by cellranger-atac (v1.2.0) with the count command line. The running parameters are used by default except for “--force-cells=”. The “--force-cell” is 10000 for liver, lung and colon, 8000 for spleen, and the rest of the organs have no restriction on this parameter. For the subsequent scATAC-seq data processing and analysis, we used the ArchR (v1.0.1) package^57^. *Macaca fascicularis* genome were constructed and annotated by createGenomeAnnotation and createGeneAnnotation function respectively. Then arrow file were created by createArrowFiles function with the default parameters. We used the addDoubletScores function to infer the doublet, filterDoublets function was used to remove the potential doublets with the “filterRatio = 1.0” parameter. ArchR project was created by ArchRProject function with the default parameters. For dimensionality reduction, we use the addIterativeLSI function in ArchR with the following parameters: “iterations = 4, clusterParams = list (resolution = c(0.2, 0.4, 0.6), sampleCells = 10000, n.start = 10, maxClusters = 6), varFeatures = 20000, dimsToUse = 1:50, scaleDims=FALSE”. Next, the Harmony package were utilized to remove the batch effect by addHarmony function^58^. AddClusters function was used to cluster cells by its default parameters. For single cell embedding, we selected the reducedDims object with harmony and used addTSNE function with the parameter “perplexity = 30” for visualization.

### Marker genes identification and cluster annotation

To identify the marker gene, gene scores were calculated when the ArchR project was created and stored in the arrow file. Then getMarkerFeatures function was used to identify the cluster-specific “expressed” genes with the default parameters. To visualize the marker genes in embedding, we used addImputeWeights function to run the MAGIC^59^ to smooth gene scores across the nearby cells. For track plot, we used the plotBrowserTrack function with the default parameters except for “tileSize = 100”.

### Peak calling and TF binding motif analysis

Before peak calling, we used addGroupCoverages function with default parameters to make pseudo-bulk replicates. Then addReproduciblePeakSet function was used with its default parameters except for “genomeSize = 2.7e09” to call accessible chromatin peaks using MACS2(v2.2.7.1)^60^. For cell type specific peak analysis, firstly, getMarkerFeatures function was applied with peak matrix to identify marker peaks. Then getMarkers function with parameter “cutOff = FDR <= 0.01 & Log2FC >= 1” was conducted to get the differential peaks. For TF motif enrichment analysis, *Macaca fascicularis* motif was downloaded from the CIS-BP database (http://cisbp.ccbr.utoronto.ca/). Then motif annotation were added to ArchR project by addMotifAnnotations function and TF motif enrichment in differential peaks were compute by the peakAnnoEnrichment function with “cutOff = FDR <= 0.1 & Log2FC >= 0.5” parameter. For motif footprint analysis, we first used getPositions function to locate relevant motifs, then getFootprints function was used to compute our interest motif footprints with its default parameters. At last footprint patterns were illustrated in plot by plotFootprints function with the following parameters: “normMethod = Subtract, smoothWindow = 10”.

### Integrative analysis of scRNA-seq and scATAC-seq data

In order to better align and integrate the scATAC-seq data, we extracted and annotated the scRNA-seq data of the 7 organs corresponding to scATAC-seq. We first used the FindTransferAnchors function from the Seurat package and aligned the data with addGeneIntegrationMatrix function in ArchR with “unconstrained integration” mode. We found in our result that most of the predicted scores > 0.5. To improve the accuracy of the predictions and better integrate the two datasets, we applied the “constrained integration” mode again to integrate the scATAC-seq and scRNA-seq data. Briefly, we annotated the scATAC-seq data with cell types based on the gene scores of scATAC-seq. Then, a restricted list were created such that gene expression similarity was calculated only in the same cell type for both scATAC-seq and scRNA-seq. For peak to gene linkage analysis, we used the addPeak2GeneLinks function to compute peak accessibility and gene expression with the parameters “corCutOff = 0.45, resolution = 1”.

## Data availability

The raw files are available from China National GeneBank (CNGB) (https://db.cngb.org/): RNA: CNP0002427; ATAC: CNP0002441. Gene counts and metadata are available at Zenodo (https://zenodo.org/): DOI: 10.5281/zenodo.5881495.

## Acknowledgements

The authors acknowledge the High Performance Computing Center of Nanjing University for providing high performance computing (HPC) resources. This work is supported by the National Natural Science Foundation of China (No. 32070656, 81872877, 82173819 and 81872876), the Nanjing University Deng Feng Scholars Program and the Priority Academic Program Development (PAPD) of Jiangsu Higher Education Institutions.

## Author contributions

D.C., Y.S. and Y.X. conceived and designed the study. Q.J., Y.W., Y.D. and X.Q. performed animal experiments. F.Y., T.Z. and W.F. performed data analyses. D.C. and J.Q. wrote the manuscript with input from X.Z. and Q.X.. All authors reviewed and approved the submitted version.

---

**Supplementary Fig. 1:**
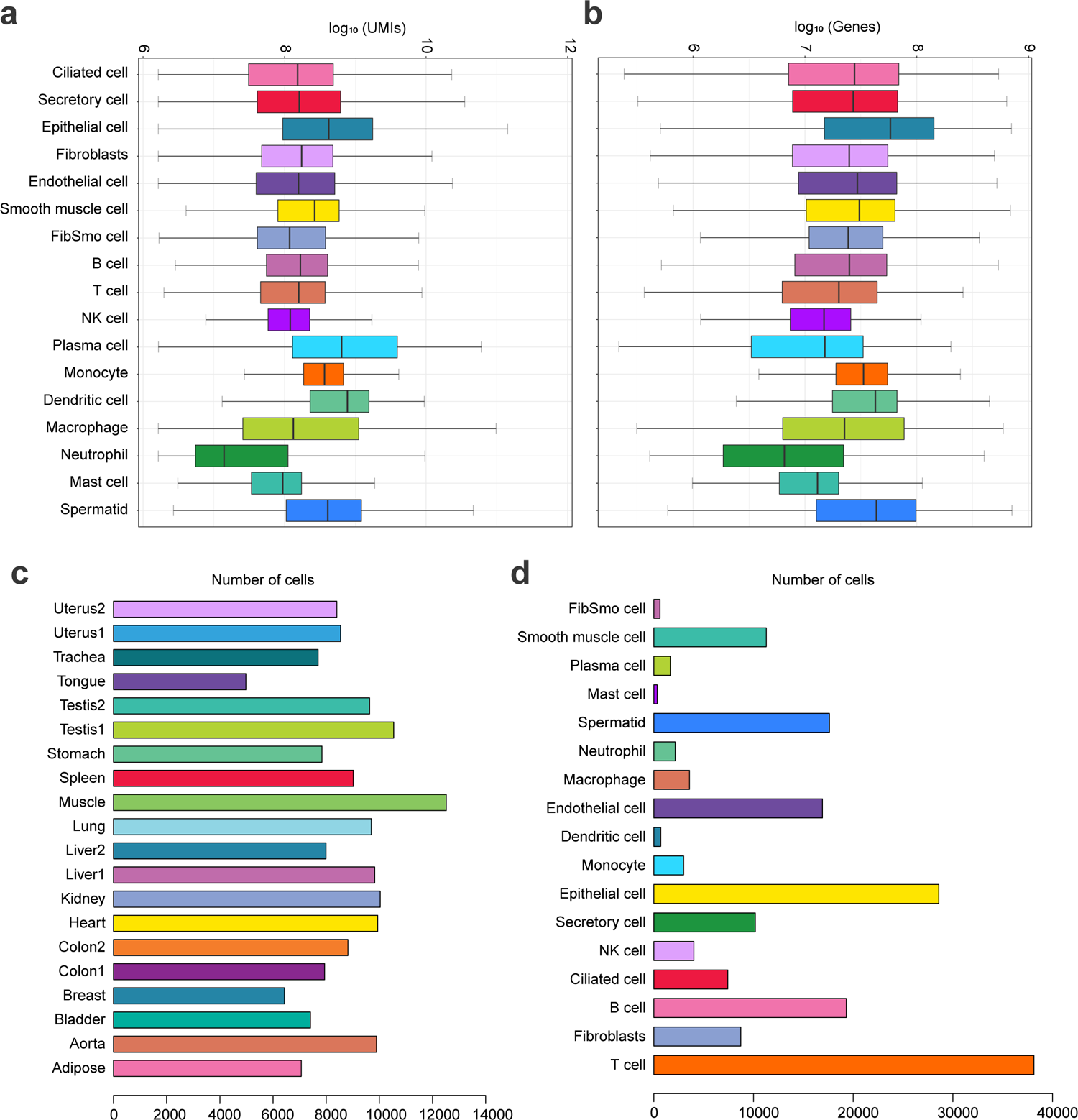
Data quality of scRNA. Related to Figure 1. **(a-b)** Box plot showing the number of UMIs and genes in major cell types respectively. **(c-d)** Bar plot showing the number of cells in major cell types and in samples.

**Supplementary Fig. 2:**
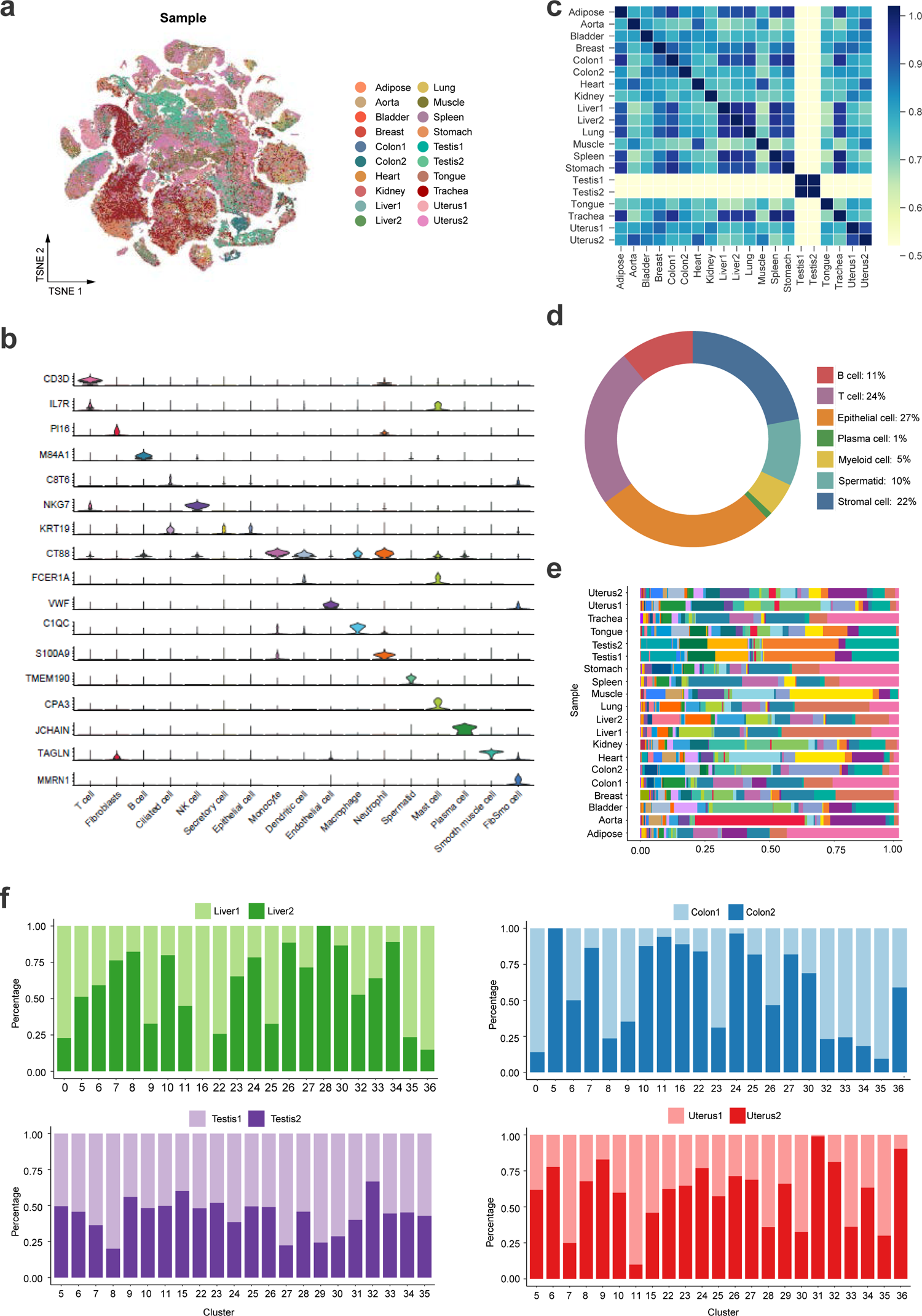
Data quality of scRNA. Related to Figure 1. **(a)** Distribution of 20 samples of cells on the tSNE. **(b)** Violin diagram showing the expression of marker genes in major cell types. **(c)** Heatmap showing correlation of highly expressed genes in 20 samples. **(d)** Circle diagram showing the cell proportions of seven cell types. **(e)** Bar plot showing the percentage of cells in each sample. **(f)** Bar Plot showing the proportion of cells in different clusters from repeated samples of liver, colon, testis and uterus.

**Supplementary Fig. 3:**
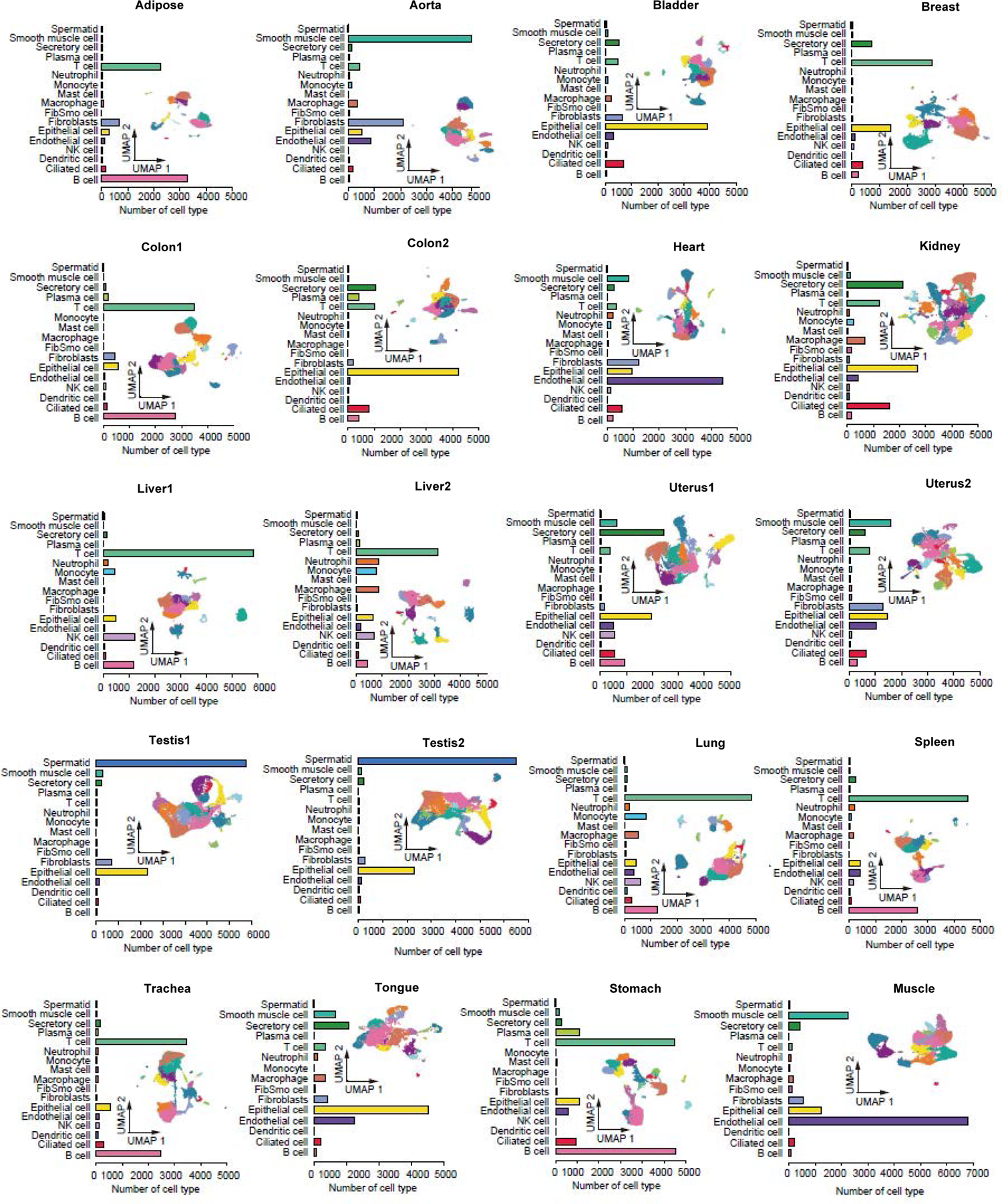
The UMAPs of all organs and bar plots of the proportion of different cell types (scRNA). Related to Figure 1. Distribution of different clusters on the UMAP in 20 samples, and bar plot shows the cell proportion of 20 samples of major cell types.

**Supplementary Fig. 4:**
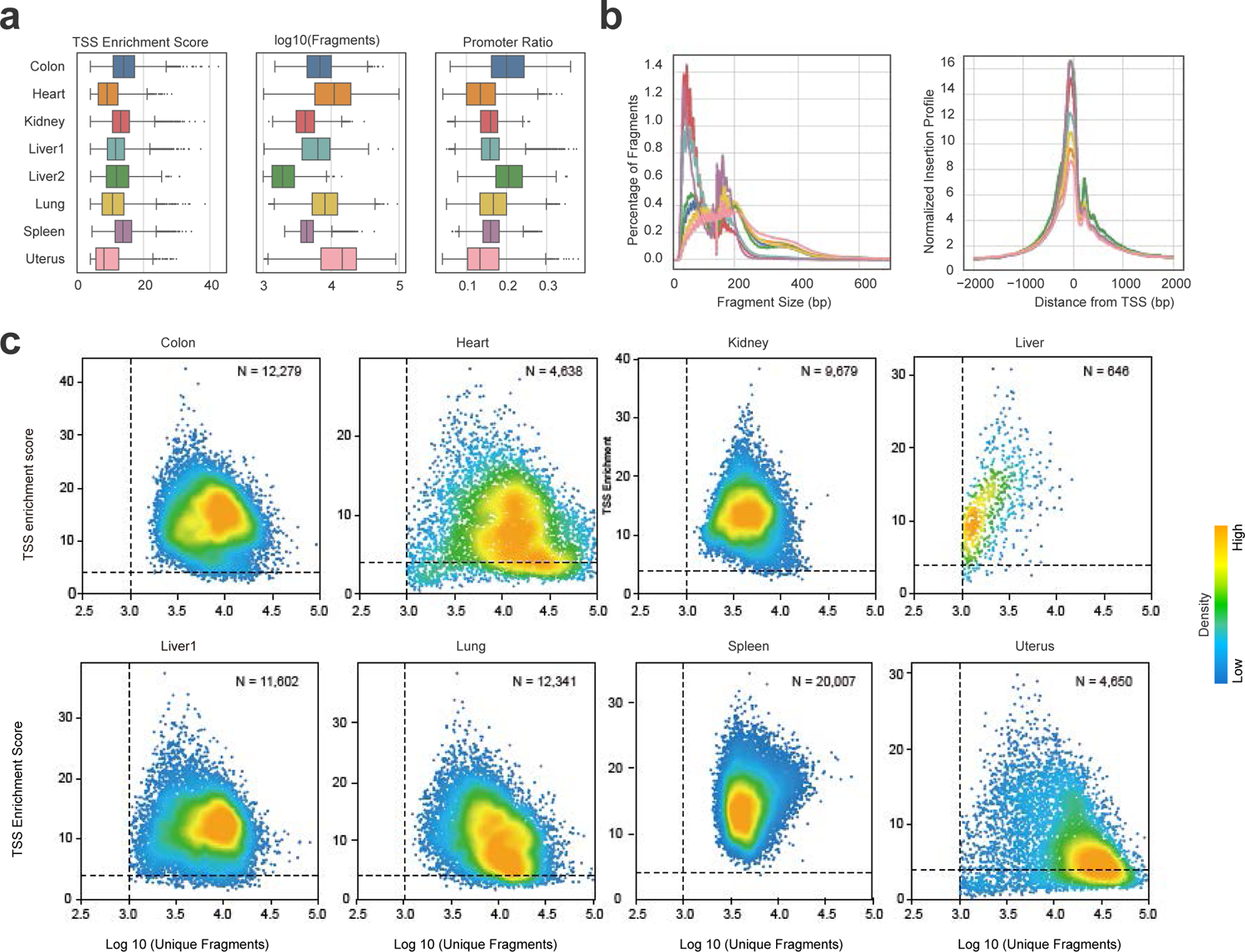
Data quality of scATAC. Related to Figure 1. **(a)** Box plot showing the distribution of TSS enrichment score, number of fragments and promoter ratio across eight samples. Each dot represents a cell. **(b)** Fragment size distributions of eight samples (left). Aggregate TSS insertion profiles centred at all TSS regions. The showing cells are passing ArchR QC thresholds for each sample (right). The line colour matches the one in a. **(c)** Scatter plot showing the TSS enrichment score vs unique nuclear fragments per cell. The colour of the dots represents the density of each point in the plot.

**Supplementary Fig. 5:**
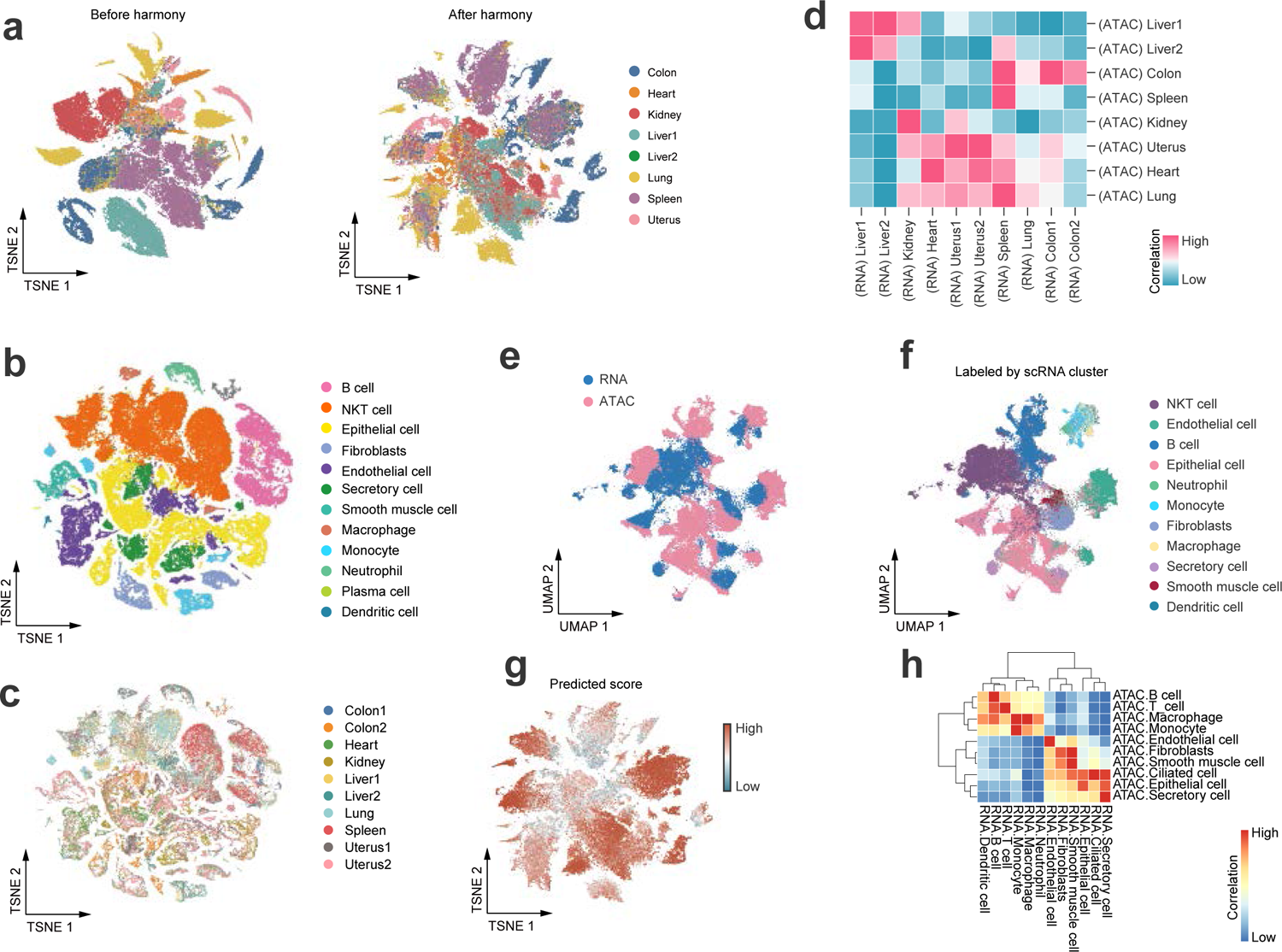
Data quality of scATAC. Related to Figure 1. **(a)** (left) TSNE of ArchR iterative LSI and (right) ArchR iterative LSI with Harmony-based batch correction for all samples. **(b)** TSNE embedding showing the subset of seven organs from scRNA-seq. Colored by cell type in Fig.1b. **(c)** TSNE embedding showing the subset of seven organs from scRNA-seq. Colored by organ. **(d)** Heatmap showing the spearman correlations between scATAC-seq and scRNA-seq samples. Correlations are calculated from the top 5,000 expressed genes for each sample. **(e)** UMAP co-embedding of scATAC-seq and scRNA-seq cell. 40,000 cells with scATAC-seq and scRNA-seq were sampled according to the percentage of cell types. **(f)** Co-embedded cells colored by scRNA-seq cell-type annotations. **(g)** Scatter plot showing the predicted score distribution for scATAC-seq and scRNAseq integration. **(h)** Heatmap showing the spearman correlations between scATAC-seq and scRNA-seq cell types. Correlations are calculated from the top 3,000 expressed genes for each cell type.

**Supplementary Fig. 6:**
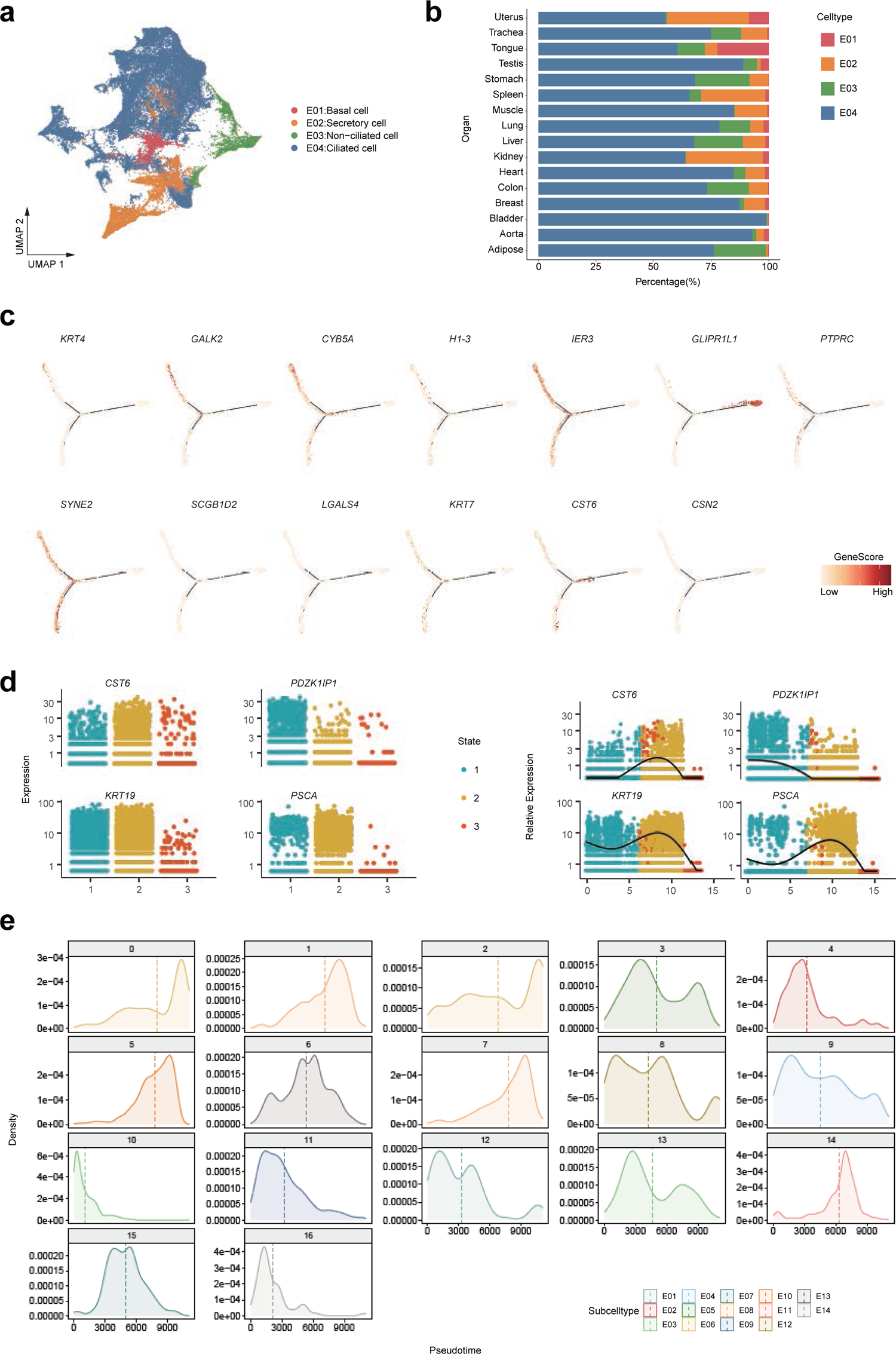
Pseudotime trajectory analysis by monocle2. Related to Figure 2. **(a)** Distribution of 4 epithelial cell subtypes on the UMAP. **(b)** Bar plot showing the percentage of cell subtypes in each organ. **(c)** Distribution of marker genes on branches during pseudotime differentiation trajectory. The shade of the color represents high or low gene scores. **(d)** Fitting curve showing the dynamic expression levels of CST6, PDZK1IP1, KRT19, PSCA in cell states. The dots represent the distribution of cells along pseudotime. **(e)** Diagrams showing the density distribution of 14 cell subtypes (16 clusters) during pseudotime differentiation trajectory.

**Supplementary Fig. 7:**
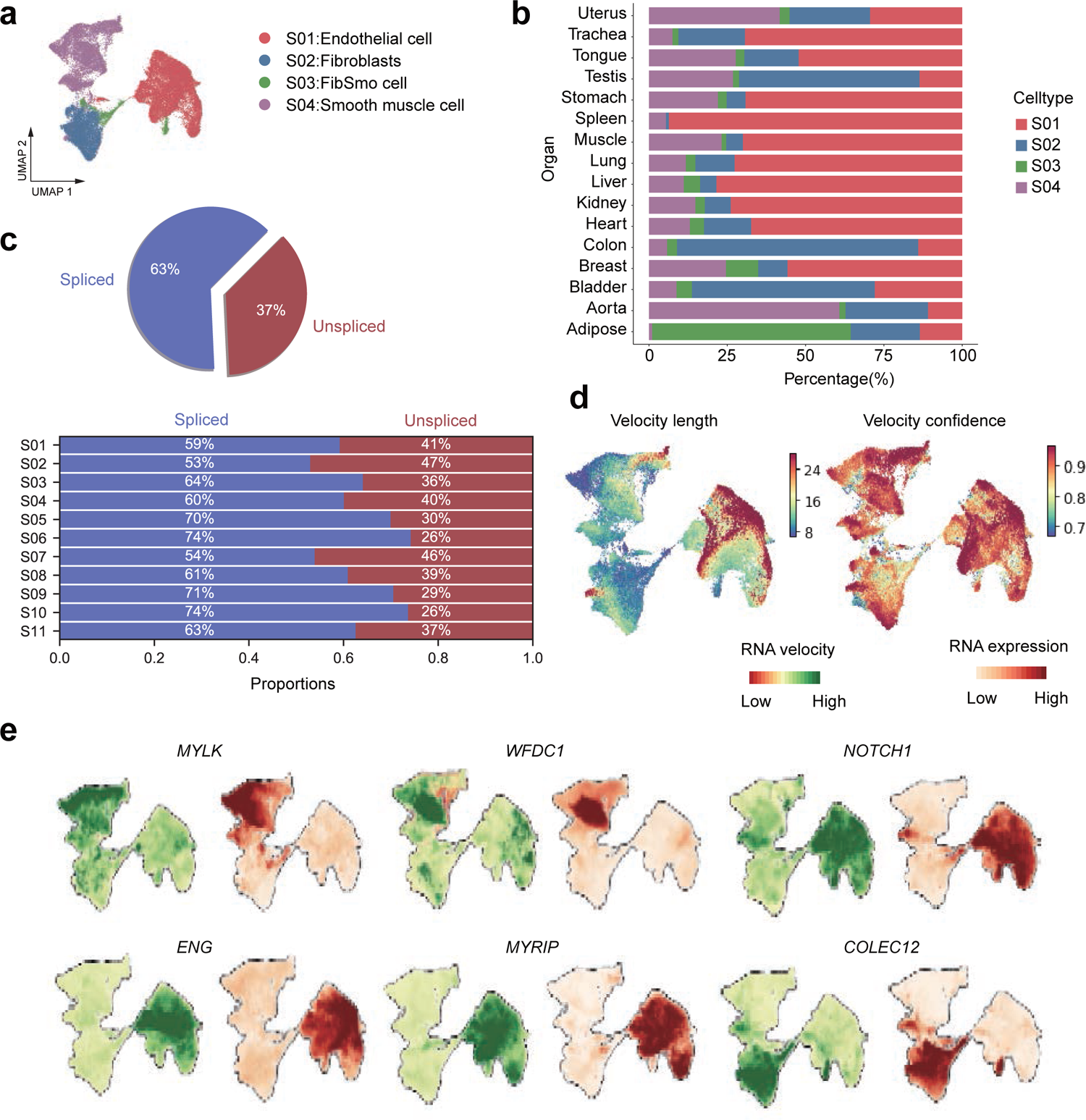
Pseudotime trajectory analysis by RNA velocity. Related to Figure 3. **(a)** Distribution of 4 stromal cell subtypes on the UMAP. **(b)** Bar plot showing the percentage of cell subtypes in each organ. **(c)** Pie chart showing the proportion of stromal cells that are spliced versus un-spliced. Bar plot showing the proportion of stromal cell subtypes that are spliced versus un-spliced. **(d)** UMAPs diagram showing the velocity length and velocity confidence of stromal cells by RNA Velocity analysis. **(e)** Distribution of marker genes on the UMAPs by running RNA Velocity.

**Supplementary Fig. 8:**
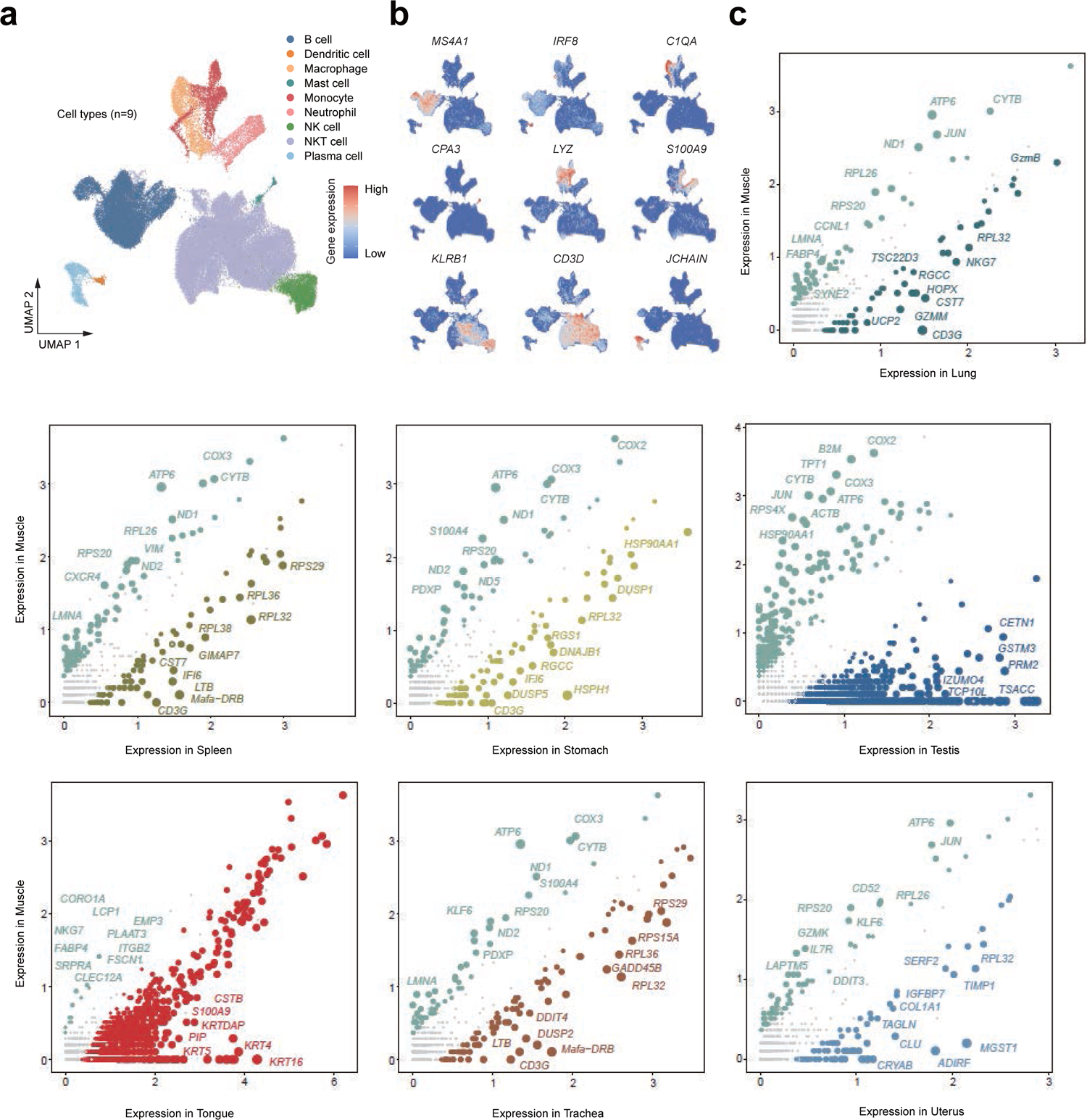
DEGs analysis of immune cell subsets. Related to Figure 4. **(a)** Distribution of 9 immune cell subtypes on the UMAP. **(b)** Feature Plot showing the expression of marker genes. **(c)** Scatter plots shows pairings of gene expression between muscle and other organs. Each point represents a DEG, and its size is proportionate to the fold change.

**Supplementary Fig. 9:**
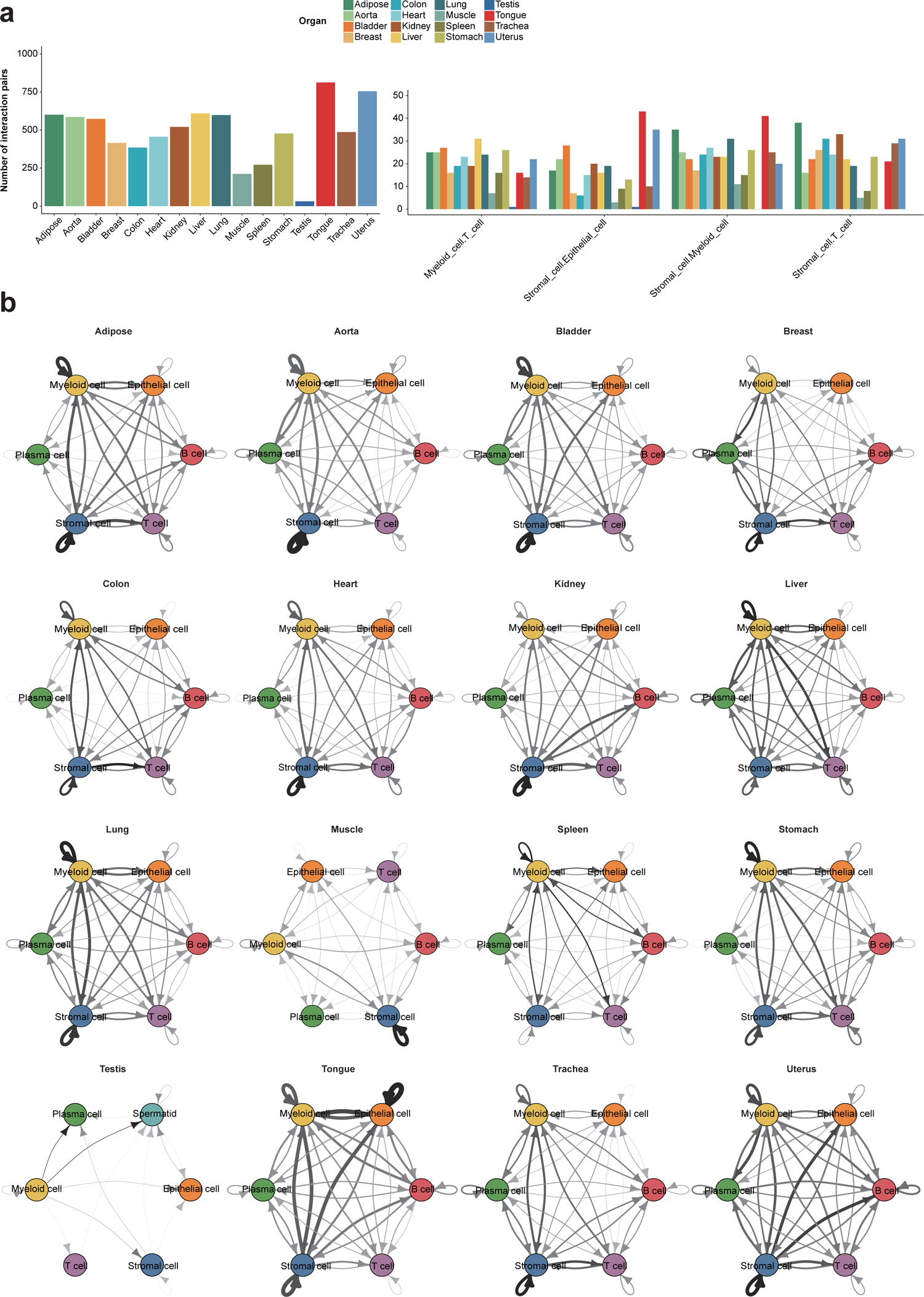
Cell-cell interactions and network diagram. Related to Figure 5. **(a)** Bar chart showing the number of interaction pairs in 16 organs (left). Bar chart showing the number of interaction pairs among four cell interaction types (Fig 6d) in 16 organs (right). **(b)** Network diagram showing cell-cell interactions among major cell types in 16 organs. Line thickness represents the strength of cell-cell interaction.

**Supplementary Fig. 10:**
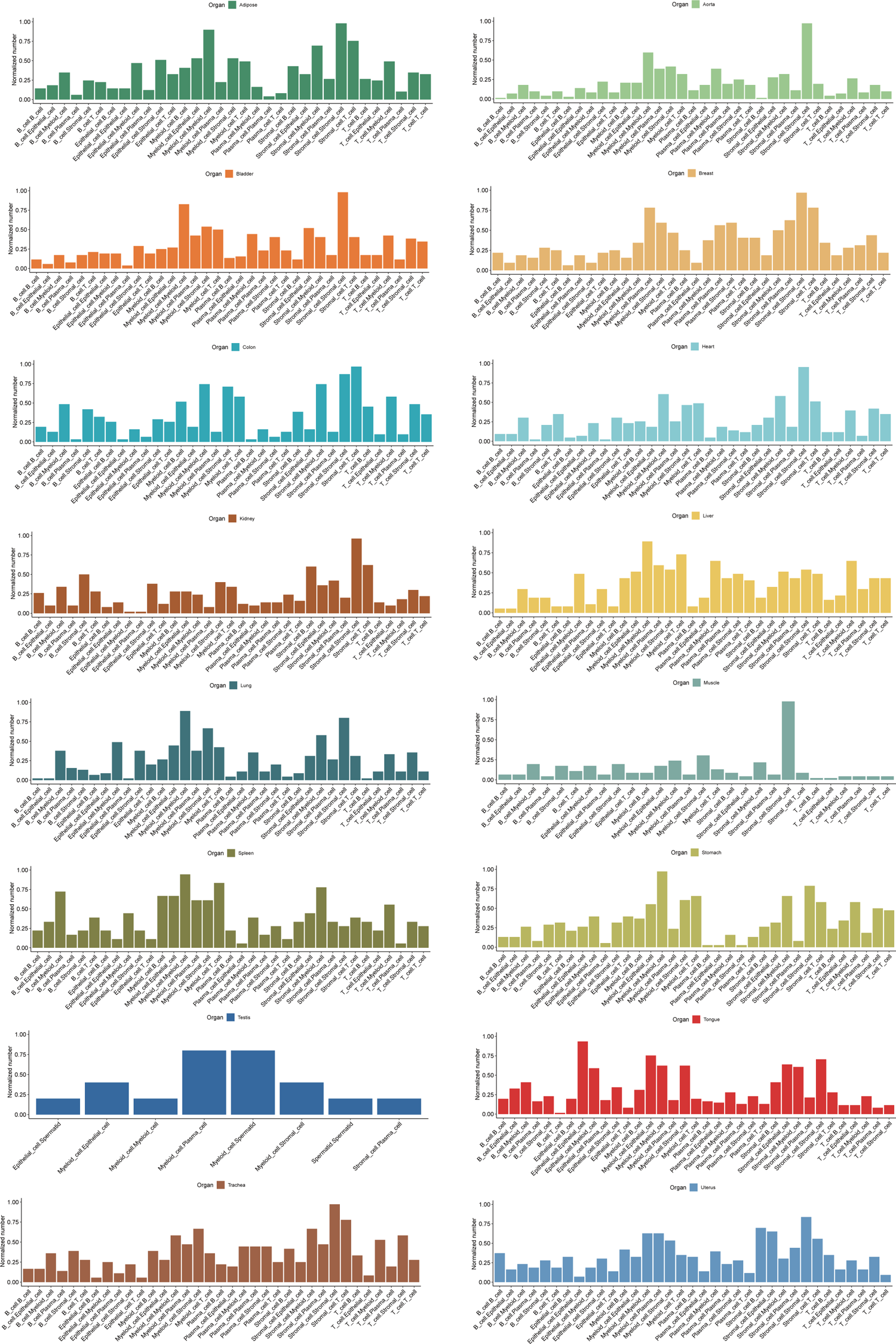
Cell-cell interactions. Related to Figure 5. Bar chart showing the normalized number of interaction pairs of different cell-cell interaction types in 16 organs.

**Supplementary Fig. 11:**
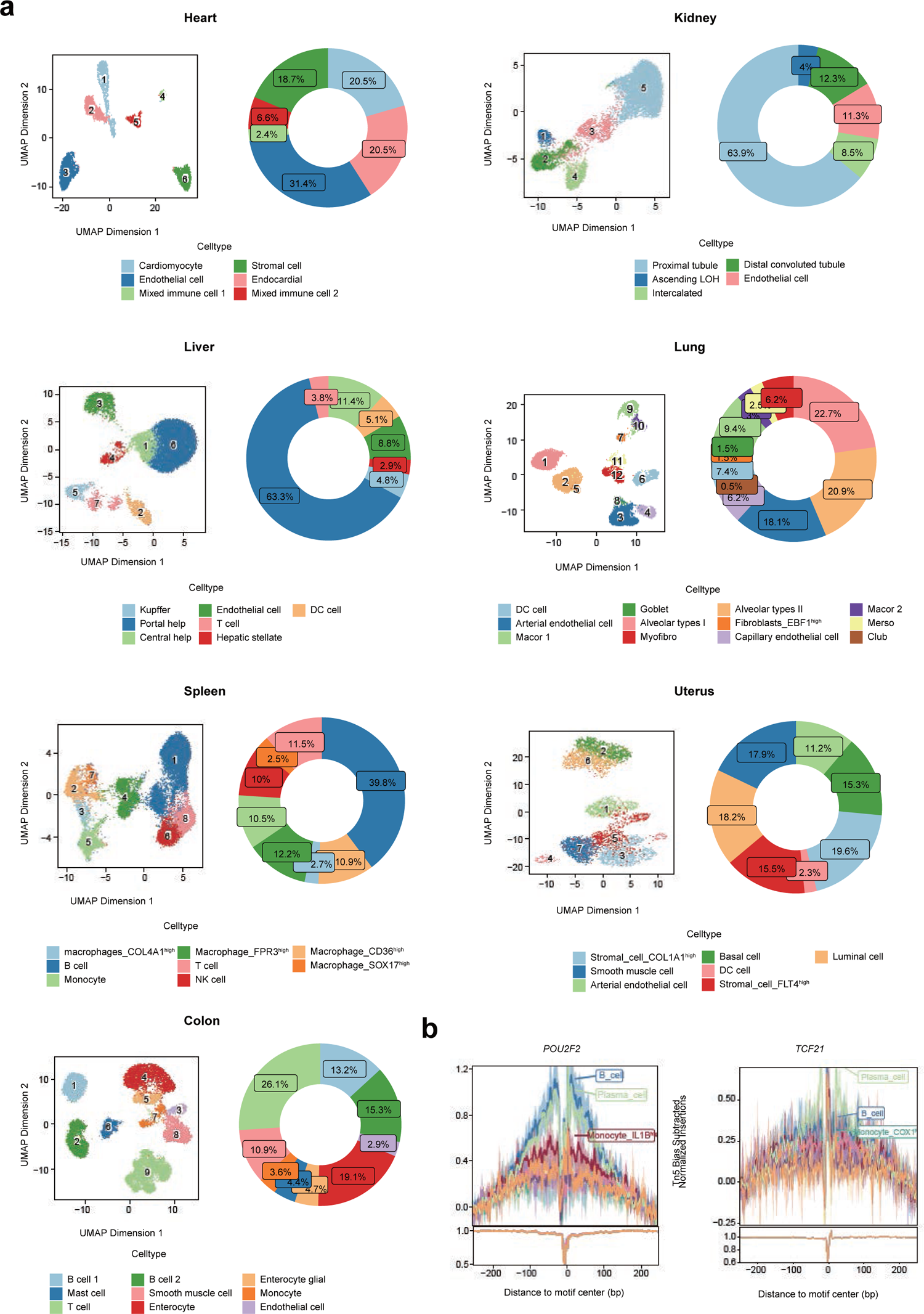
The UMAPs of all organs and Donut charts of the proportion of different cell types (scATAC). Related to Figure 6. **(a)** Distribution of different clusters on the UMAP in 7 organs (scATAC-seq), and donut chart shows the cell proportion of 7 organs of cell types. **(b)** Tn5 bias-adjusted TF footprint analysis of the POU2F2 and TCF21 transcription factors.

**Supplementary Fig. 12:**
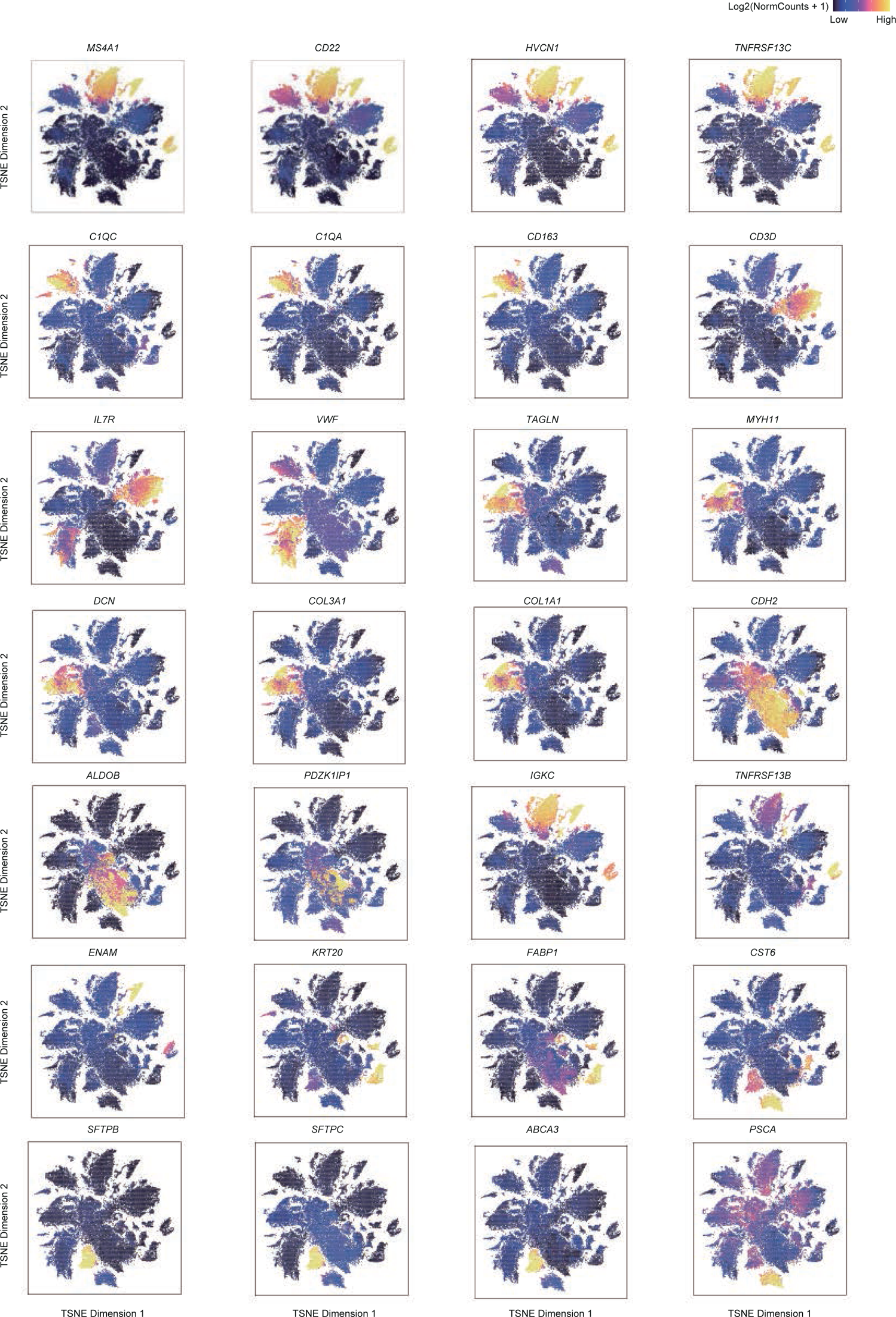
The TSNEs of marker genes expression (scATAC). Related to Figure 6. Distribution of marker genes in cell types such as B cells (MS4A1, CD22, HVCN1, TNFRSF13C, IGKC, TNFRSF13B, ENAM), macrophages (C1QC, C1QA, CD163), T cells (CD3D, IL7R), endothelial cells (VWF), fibroblasts (TAGLN, MYH11, DCN, COL3A1, COL1A1), secretory cells (CDH2, ALDOB, PDZK1IP1), epithelial cells (KRT20, FABP1) and ciliated cells (CST6, SFTPB, SFTPC, ABCA3, PSCA) on the TSNEs.

**Supplementary Fig. 13:**
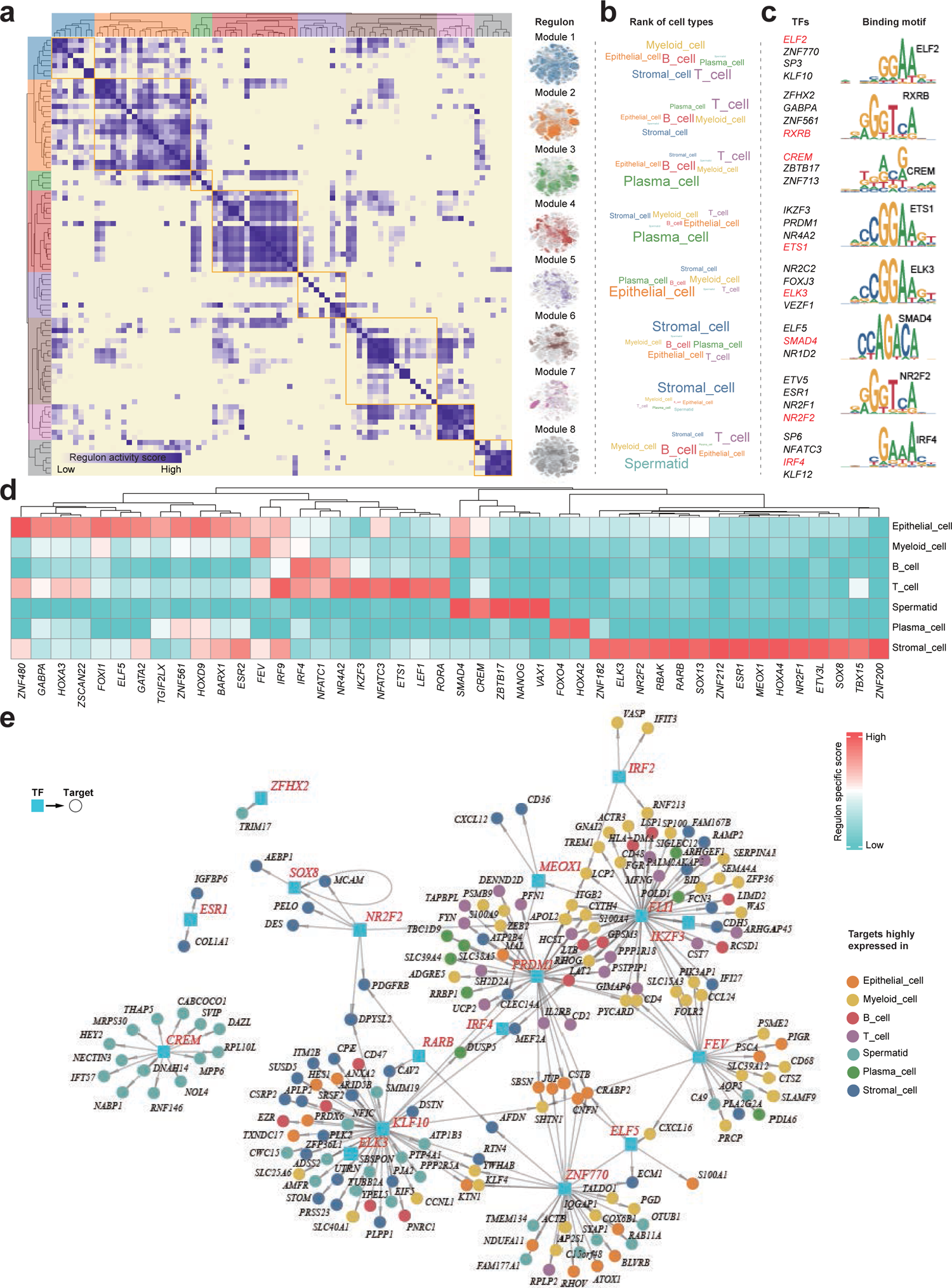
Transcriptional regulatory network of cells from cynomolgus monkey. Related to Figure 7. **(a)** Identification of regulon modules using SCENIC. Heatmap (left) shows the similarity of different regulons (n=86) based on the AUCell score. Eight regulon modules were identified based on regulon similarity. UMAPs (right) illustrate the average AUCell score distribution for different regulon modules (in different colors). **(b)** Wordcloud plots shows enrichment of cell types in different regulon modules. **(c)** Representative transcription factors and corresponding binding motifs in different regulon modules. **(d)** Heatmap showing transcription factors enriched in different cell types. Color depth represents the level of regulon-specific score. **(e)** Integrated gene-regulatory networks of the regulons from (a). Regulon-associated TFs are highlighted in blue rectangles and target genes in circles. Target genes (in circles) are colored according to their highly expressed cell types.

**Supplementary Fig. 14:**
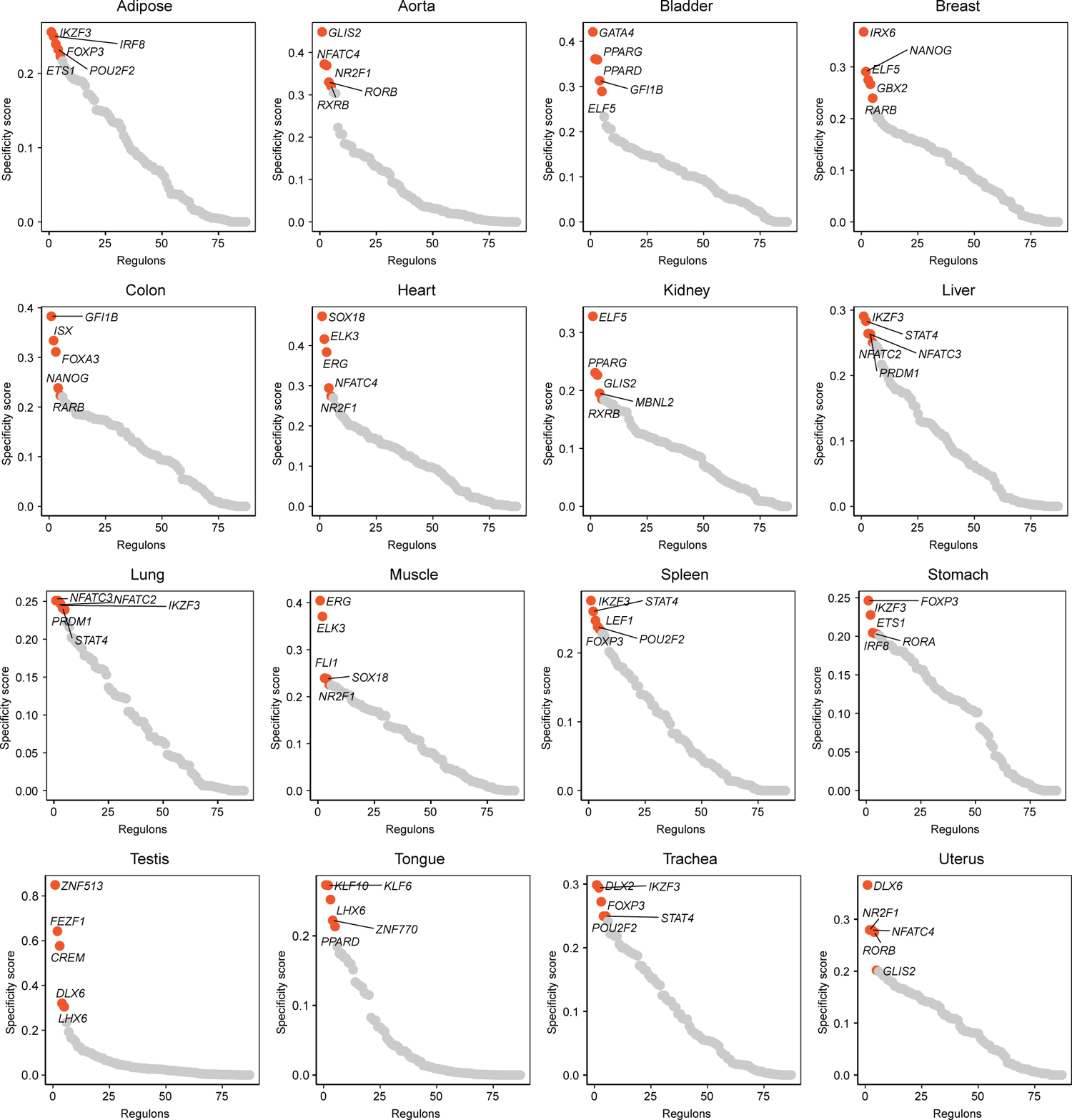
Regulons and modules. Related to Figure 7. Scatter plot showing top regulons of 16 organs ordered by regulon specific score and red dots represent the highest activity.

**Supplementary Fig. 15:**
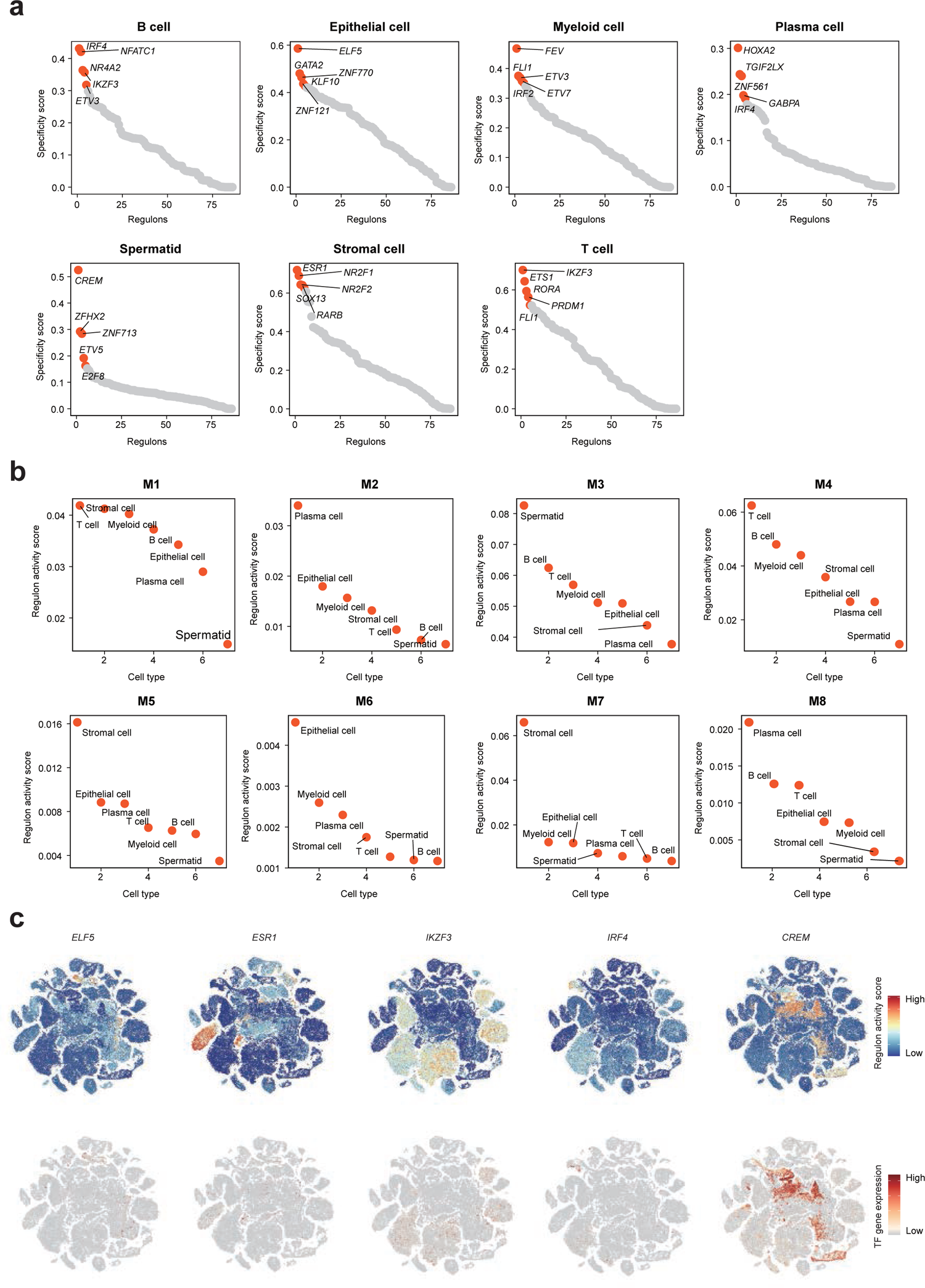
Regulons and modules. Related to Figure 7. **(a)** Scatter plot showing top regulons of 7 major cell types ordered by regulon specific score and red dots represent the highest activity. **(b)** Scatter plot showing major cell types of 8 regulon modules ordered by regulon activity score. **(c)** TSNEs displaying the AUCell score distribution for the selected regulons (above) and gene expression patterns for the corresponding TFs (below).

**Supplementary Fig. 16:**
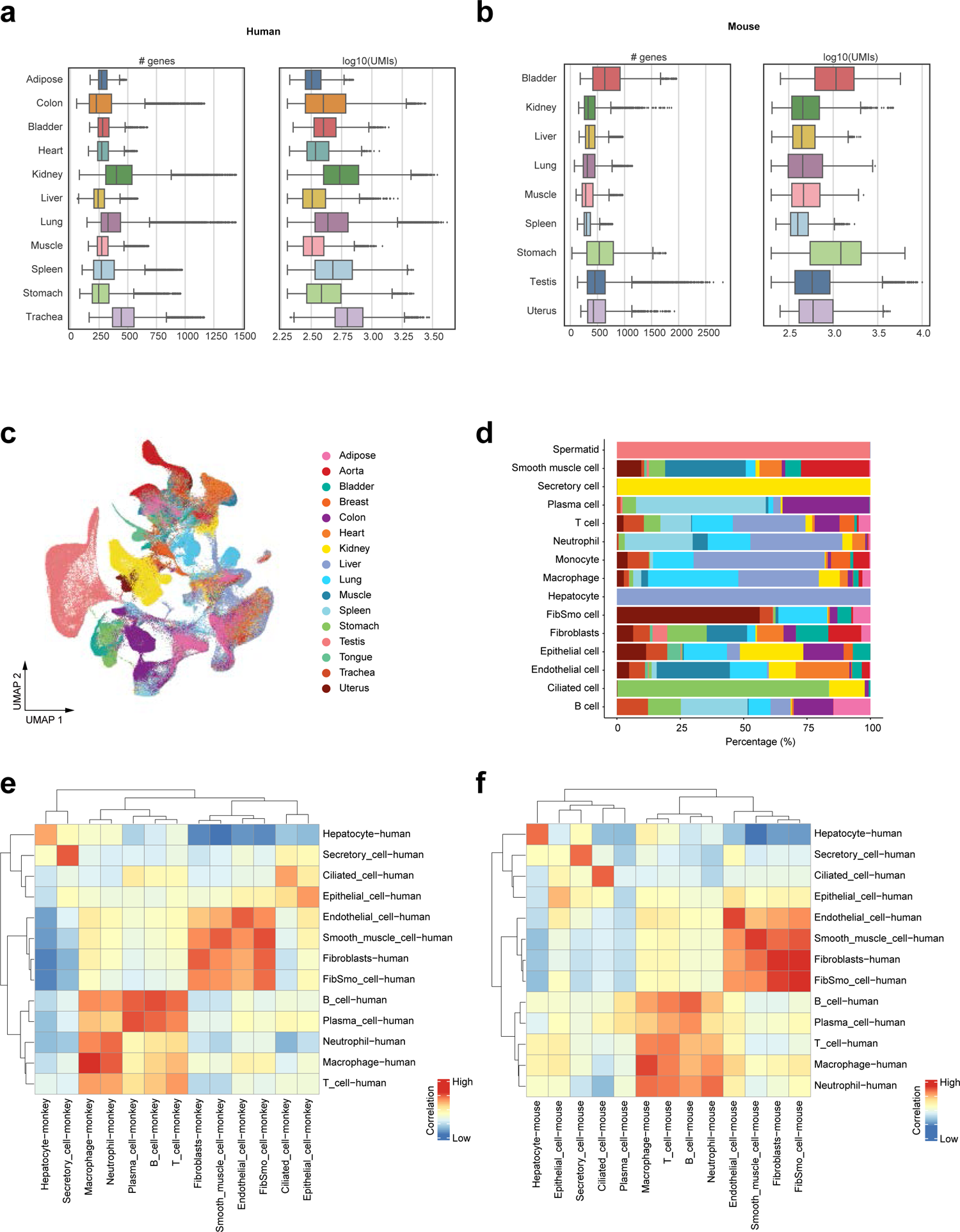
Data quality across species. Related to Figure 8. **(a)** Box plot showing the number of genes and UMIs in human organs. **(b)** Box plot showing the number of genes and UMIs in mouse organs. **(c)** Distribution of cynomolgus monkey, human and mouse cells by organs on the UMAP. **(d)** Bar plot showing the percentage of cell types in each organ. **(e)** Heatmap showing correlation of top gene expression in cell types in mouse and human (Spearman method). **(f)** Heatmap showing correlation of top genes expression in cell types cynomolgus monkey, and human (Spearman method).

